# Telomerase RNA regulates the epigenome primed for human lineage commitment

**DOI:** 10.64898/2026.07.03.736284

**Authors:** Jie Li, Peng Su, Mingqi Gao, Chang Liu, Niannian Li, Guofeng Feng, Yongqin Yu, Zhifei Chen, Guoxing Yin, Xiaoying Ye, Jiangtao Lu, Ziyi Jin, Zhengmao Zhu, Haiying Liu, Hua Wang, Lin Liu

## Abstract

Telomerase RNA (*TERC*) is known as the essential template for telomere elongation. Here, we report an unexpected role for *TERC* in regulating chromatin accessibility, which is primed for determining the cell fate of human embryonic stem cells (hESCs). *TERC*-deficient hESCs retain critical markers for pluripotency but fail to undergo lineage differentiation as shown by standard *in vivo* teratoma formation as well as *in vitro* differentiation assays, which is consistent with repressed transcription during differentiation into the three germ lineages. Notably, transient re-introduction of *TERC* into *TERC*-deficient hESCs rescued lineage differentiation capacity without restoring telomere length. *TERC* binds to the promoters and enhancers of developmental genes marked by H3K27ac to maintain an open chromatin state. Loss of *TERC* reduces H3K27ac deposition and decreases chromatin accessibility through the remodelling of three-dimensional genome organization, including TAD boundary insulation and compartment switching. Collectively, our findings reveal that *TERC* is a chromatin-associated noncoding RNA that regulates the epigenomic architecture that governs cell fate for lineage commitment during development.

## Introduction

Telomerase is a ribonucleoprotein complex whose catalytic core consists of telomerase reverse transcriptase (*TERT*) and its essential RNA component *(TERC*), which provides the template for telomeric repeat synthesis at chromosome ends to maintain telomeres and genome stability^1–6^. Germline mutations that disrupt *TERC* and its stability and reduce telomerase function are associated with a spectrum of telomere biology disorders (TBDs), including dyskeratosis congenita, pulmonary fibrosis, aplastic anemia, and immunodeficiency^7–15^. In addition to its canonical role in templating telomeric DNA synthesis to preserve genomic stability, *TERC* also performs noncanonical, template-independent functions^16,17^. This functional dichotomy is highlighted by the fact that complete *Terc* knockout mice develop and survive to adulthood normally, with defects manifesting only in late generations upon critical telomere shortening, which is prone to tumorigenesis and aging^18–22^. These observations suggest that the shortest telomeres impair development but that *Terc* itself is dispensable for lineage differentiation and development in mice. Notably, mouse *Terc* shares only 65% sequence identity with human *TERC*^23^, underscoring potential interspecies functional differences.

The study of *TERC* function in human development is inherently limited; thus, human embryonic stem cells (hESCs), which can differentiate into all three germ layer lineages^24–26^, constitute an essential complementary model. While *Terc*-deficient mouse ESCs retained the ability to differentiate into all three germ layers according to a teratoma formation assay, reduced differentiation potential was observed only in the late G3 and G4 generations of *Terc-*deficient ESCs with critically short or lost telomeres^27^. These results support the notion that the shortest telomeres reduce differentiation and that *Terc* deficiency itself does not affect cell differentiation in mice^28–31^. The ESC *in vitro* differentiation model is consistent with the *in vivo* development of the mouse model described above. Additionally, *Terc* is dispensable for iPSC derivation in mice^32^, whereas human iPSCs cannot be derived without *TERC*^33^. These results suggest fundamental differences in *TERC* function during development between species^7^.

We therefore tested the hypothesis that human *TERC* may have additional functions in lineage differentiation during human development. Here, we report that *TERC,* as a chromatin-associated regulatory gene, governs human lineage differentiation into all three germ layers.

## Results

### *TERC* is required for lineage differentiation of hESCs

We generated *TERC* knockout (KO) hESCs in two independent cell lines, WA26 and H1, using CRISPR-Cas9 for subsequent phenotypic analysis, rescue validation, and multi-omics profiling (Fig. 1a and Extended Data Fig. 1a–c). Compared with those of WT hESCs, the colonies of *TERC* KO hESCs were smaller and less compact (Fig. 1b). Consistently, *TERC* knockout resulted in telomerase-negative hESCs, as measured by a telomere repeat amplification (TRAP) assay (Extended Data Fig. 1d, e). *TERC* KO hESCs presented decreased telomere length, as determined by telomere restriction fragment (TRF) Southern blotting, telomere quantitative fluorescence *in situ* hybridization (Q-FISH) and the T/S ratio (Extended Data Fig. 1f–h). The protein levels of key pluripotency markers (OCT4, NANOG and SSEA4) did not significantly change following *TERC* knockout (Fig. 1c, d, and Extended Data Fig. 1i). Thus, *TERC* is essential for telomere homeostasis but does not affect the expression of key pluripotency factors.

**Fig. 1.**
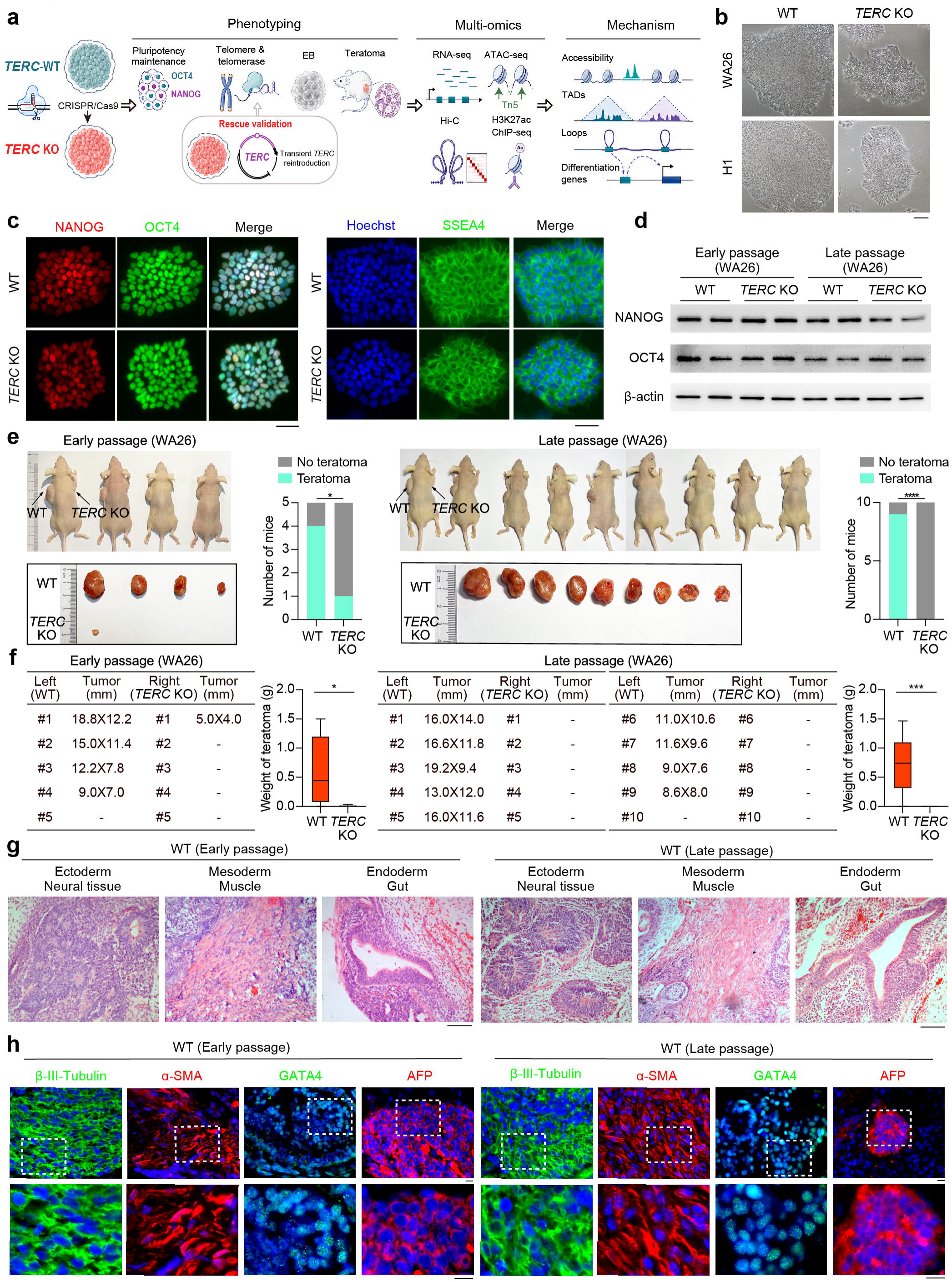
*TERC* is dispensable for pluripotency maintenance but is required for the differentiation potential of hESCs. **a,** Overview of the experimental design. WT and *TERC* KO hESCs were used for phenotypic analysis, rescue validation, and multi-omics profiling, including RNA-seq, ATAC-seq, H3K27ac ChIP-seq, and Hi-C, to investigate the functions of *TERC*. **b,** Representative brightfield images showing the morphology of WT and *TERC* KO hESCs (WA26 and H1). Scale bar = 100 μm. **c,** Representative immunofluorescence images of the pluripotency markers OCT4, NANOG and SSEA4 in *TERC* KO and WT hESCs (WA26), and nuclei were counterstained with Hoechst (blue). Scale bars, 50 μm. **d,** Western blot analysis of pluripotent factors (OCT4 and NANOG) in *TERC* KO and WT hESCs (WA26; early and late passages). n=2 biological replicates. **e,** Teratoma formation assay using *TERC* KO and WT hESCs (WA26) at early (left) and late (right) passages in immunodeficient NU/NU mice. The black arrows indicate the injection sites of *TERC* KO and WT mice. n=5/10 biological replicates, and *P* values were calculated via the chi-square test. **f,** Representative images and quantification of the size and weight of the teratomas 8 weeks after subcutaneous injection into the left or right of NU/NU mice with the indicated hESCs (WA26). The weights of the teratomas are shown in box plots (n=5/10 biological replicates), and *P* values were calculated via the Mann‒Whitney test. **g,** Hematoxylin and eosin (H&E) staining of teratomas derived from WT hESCs (WA26) at early (left) and late passages (right), showing tissues from the ectoderm (neural), mesoderm (muscle), and endoderm (gut). Scale bar, 100 μm. **h,** Immunofluorescence staining for markers of three germ layers from WT hESCs (WA26) at early- (left) and late-passage (right)-derived teratomas, including β-III-tubulin (ectoderm), α-SMA (mesoderm), GATA4 and AFP (endoderm). Scale bar, 10 μm.

To evaluate the effect of *TERC* depletion on the *in vivo* differentiation potential of hESCs, we performed a standard teratoma formation assay using immunodeficient mice. Strikingly, compared with WT hESCs, early- and late-passage *TERC* KO hESCs exhibited severely impaired teratoma formation, generating only small teratomas or failing to form teratomas (Fig. 1e, f, and Extended Data Fig. 1j, k), indicating that the differentiation potential of *TERC* KO hESCs was compromised. While WT hESC-derived teratomas contained well-differentiated tissues representing all three germ layers (including neural tissue, muscle, and gut epithelium), *TERC* KO hESC-derived teratomas showed a complete absence of intact or properly organized germ layer structures (Fig. 1g and Extended Data Fig. 1l), and immunofluorescence analysis further confirmed this observation, revealing appropriate expression of characteristic markers associated with the neuroectoderm, mesoderm, and endoderm lineages in teratomas derived from WT hESCs (Fig. 1h and Extended Data Fig. 1m). Taken together, these findings indicate that *TERC* deficiency impairs differentiation in hESCs.

### *TERC* deficiency disrupts developmental gene expression programs

To better understand the molecular function of *TERC* in hESCs, we performed RNA-seq analysis of WT and *TERC* KO hESCs. *TERC* knockout resulted in significant changes in the global transcriptome of hESCs (Extended Data Fig. 2a), with 2059 upregulated genes and 1265 downregulated genes in WT hESCs compared with those in *TERC* KO hESCs (Fig. 2a). In agreement with the observed changes in hESC morphology, stem cell proliferation was slightly reduced following *TERC* knockout (Extended Data Fig. 2b, c). The expression of most pluripotency-related genes, such as the core pluripotency factors *POU5F1* and *NANOG* (Extended Data Fig. 2d), did not change significantly after *TERC* knockout, which was consistent with the changes in protein levels (Fig. 1c, d and Extended Data Fig. 1i). Notably, we found that *TERC* knockout significantly suppressed key signalling pathways, including the MAPK and PI3K-AKT pathways (Fig. 2b, c), which are essential for hESC differentiation^34–36^. Moreover, *TERC* knockout resulted in decreased expression of genes related to many developmental processes, such as angiogenesis, brain development, nervous system development and skeletal system development (Fig. 2d), and these findings were also observed in an independent hESC line (H1 hESCs) (Extended Data Fig. 2e–i). This prompted us to explore the changes in the expression of genes related to the three germ layers. Interestingly, many endodermal, mesodermal and ectodermal-specific genes were markedly downregulated in *TERC* KO hESCs (Fig. 2e). These data indicate that *TERC* is essential for the expression of development-related genes.

**Fig. 2.**
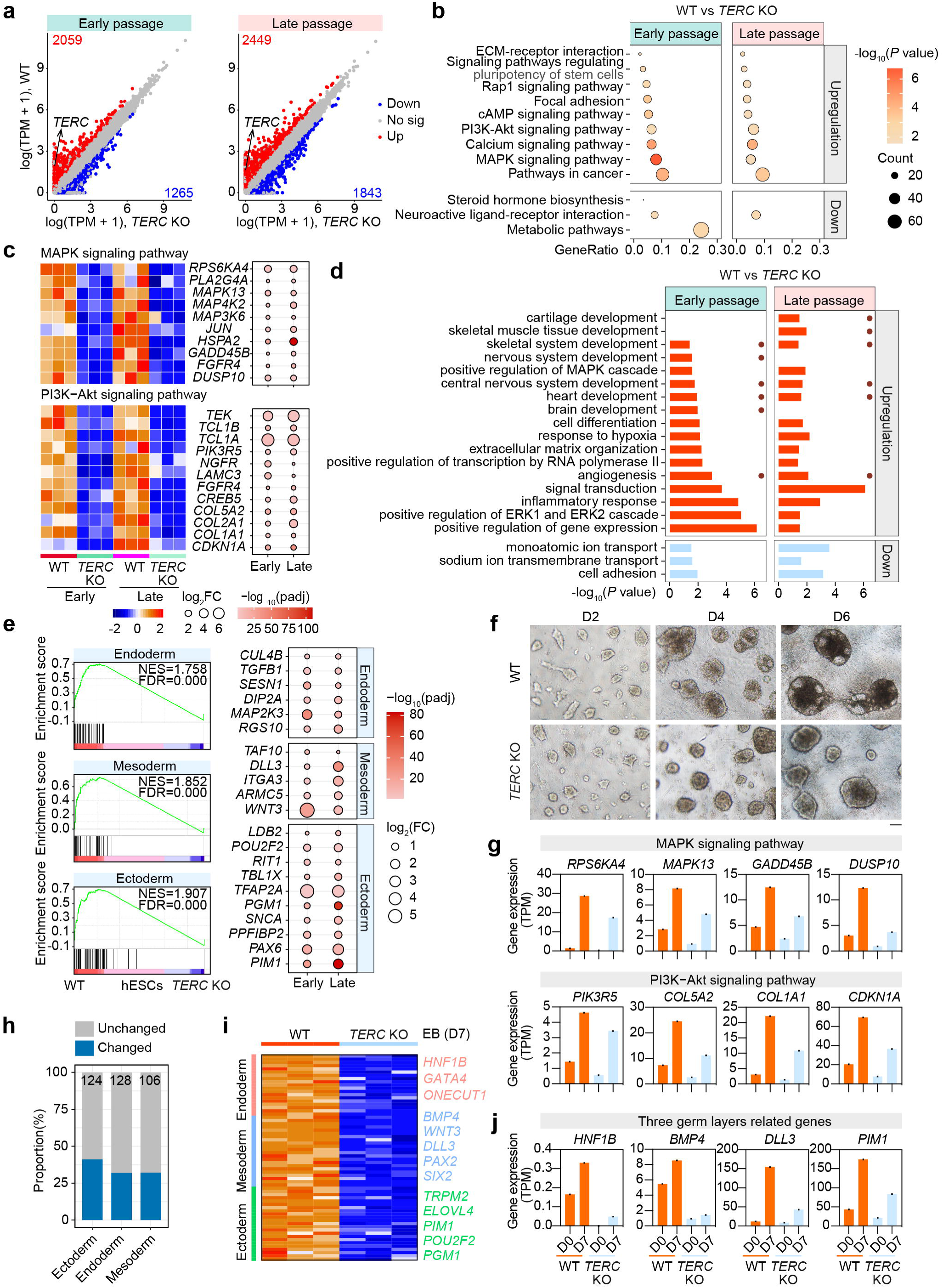
TERC regulates the transcriptional program of lineage differentiation. **a,** Scatterplot comparing the gene expression levels of WT and *TERC* KO hESCs (WA26; early and late passages). The red and blue dots represent genes whose expression was upregulated and downregulated, respectively, in WT hESCs compared with *TERC* KO hESCs. **b,** KEGG analysis showing key signalling pathways enriched in up- or downregulated genes in WT hESCs (WA26; early and late passages) compared with *TERC* KO hESCs. The horizontal axis represents the gene ratio, the size of the circles represents the gene count, and the color depth represents the -log10-transformed *P* value. **c,** Heatmap depicting the expression of MAPK and PI3K-Akt signalling pathway-related genes, as revealed by RNA-seq analysis. **d,** GO analysis showing representative terms enriched with up- or downregulated genes in WT hESCs (WA26; early and late passages). Red and blue represent GO terms enriched with upregulated and downregulated genes, respectively. **e,** GSEA showing enrichment of lineage-specific gene sets (endoderm, mesoderm, and ectoderm) in WT hESCs. Red, upregulated genes in WT hESCs; blue, downregulated genes in WT hESCs; NES, normalized enrichment score; FDR, false discovery rate. Bubble plots showing the log2(FC, fold change) and −log10(padj, adjusted *P* value) values of three key germ layer-related genes between groups (WT vs *TERC* KO). **f,** Representative images of EBs derived from WT and *TERC* KO hESCs at days 2, 4, and 6 of *in vitro* differentiation. Scale bar, 100 μm. **g,** Changes in the expression levels of MAPK and PI3K-Akt signalling pathway-related genes during *in vivo* differentiation following *TERC* knockout. **h,** The bar graph shows the percentage of germ layer-related genes whose expression was altered in *TERC* KO EBs compared with that in WT EBs. Blue indicates significantly changed genes, gray represents unchanged genes, and the total number of genes associated with each germ layer is indicated above the corresponding bars. **i,** Heatmap showing the significantly downregulated expression of three germ layer-related genes in *TERC* KO EBs relative to WT EBs. Representative marker genes are labelled. **j,** Changes in the expression levels of three germ layer-related genes during *in vivo* differentiation after *TERC* knockout.

We subsequently performed *in vitro* differentiation assays involving the formation of embryoid bodies (EBs) and reported that *TERC* deficiency led to smaller, structurally disorganized EBs with compromised differentiation potential, which aligns with the RNA-seq findings of dysregulated developmental genes (Fig. 2f). Therefore, we collected day 7 EBs for further RNA-seq analysis and found that *TERC* KO resulted in significant changes in the transcriptome of EBs (Extended Data Fig. 3a), including 2498 genes whose expression was upregulated and 2080 genes whose expression was downregulated in the WT control (Extended Data Fig. 3b). Enrichment analysis further revealed that genes upregulated in WT controls were associated primarily with key signalling pathways, including the MAPK and PI3K-AKT pathways (Extended Data Fig. 3c). Notably, these pathways were also significantly enriched in WT hESCs (Fig. 2b), and key genes of these two signalling pathways were not adequately activated during the differentiation process (Fig. 2g), suggesting that *TERC* is critical for the proper activation of signalling pathways such as the MAPK and PI3K-Akt pathways during differentiation. Importantly, genes upregulated in WT EBs were significantly enriched in key developmental processes, particularly those involved in angiogenesis and nervous system development (Extended Data Fig. 3d). The appropriate and precise expression of key genes for the differentiation of the three germ layers is a prerequisite for subsequent developmental processes^37^. However, *TERC* deficiency disrupted the expression of more than 30% of the genes associated with each germ layer (Fig. 2h, i). Furthermore, the deletion of *TERC* abrogated the appropriate activation of these genes during differentiation (Fig. 2j), which may have culminated in the failure of differentiation.

In short, transcriptional dysregulation in *TERC* KO EBs is consistent with impaired differentiation capacity, highlighting a critical role for *TERC* in regulating the proper expression of key developmental genes across all three germ layers.

### Transient re-introduction of *TERC* rescues the differentiation defects of *TERC* KO hESCs

To determine whether *TERC* directly regulates hESC differentiation independent of its telomerase function, we transiently re-introduced *TERC* into *TERC* KO hESCs using an overexpression construct. This transient approach was chosen to generate stable cell lines to minimize the potential role of *TERC* in stem cell proliferation. After 48 hours of nucleofection and puromycin selection, *TERC*-overexpressing hESCs restored *TERC* RNA expression to 45% of the WT level, as determined by qPCR (Extended Data Fig. 4a), which was accompanied by partial restoration of telomerase activity and minimal telomere elongation (Fig. 3a–c). Compared with those in WT hESCs, the telomere length in these rescued cells remained significantly shorter. Furthermore, *TERC* reintroduction did not alter the expression levels of the core pluripotency factors OCT4 and NANOG (Fig. 3d and Extended Data Fig. 4b). Notably, the gene expression profile of rescued *TERC* hESCs closely resembled that of WT hESCs at the transcriptome-wide level (Fig. 3e). Further differential expression and enrichment analyses revealed that many genes, functions and signalling pathways could be rescued by *TERC* re-introduction (Fig. 3 f, g and Extended Data Fig. 4c).

**Fig. 3.**
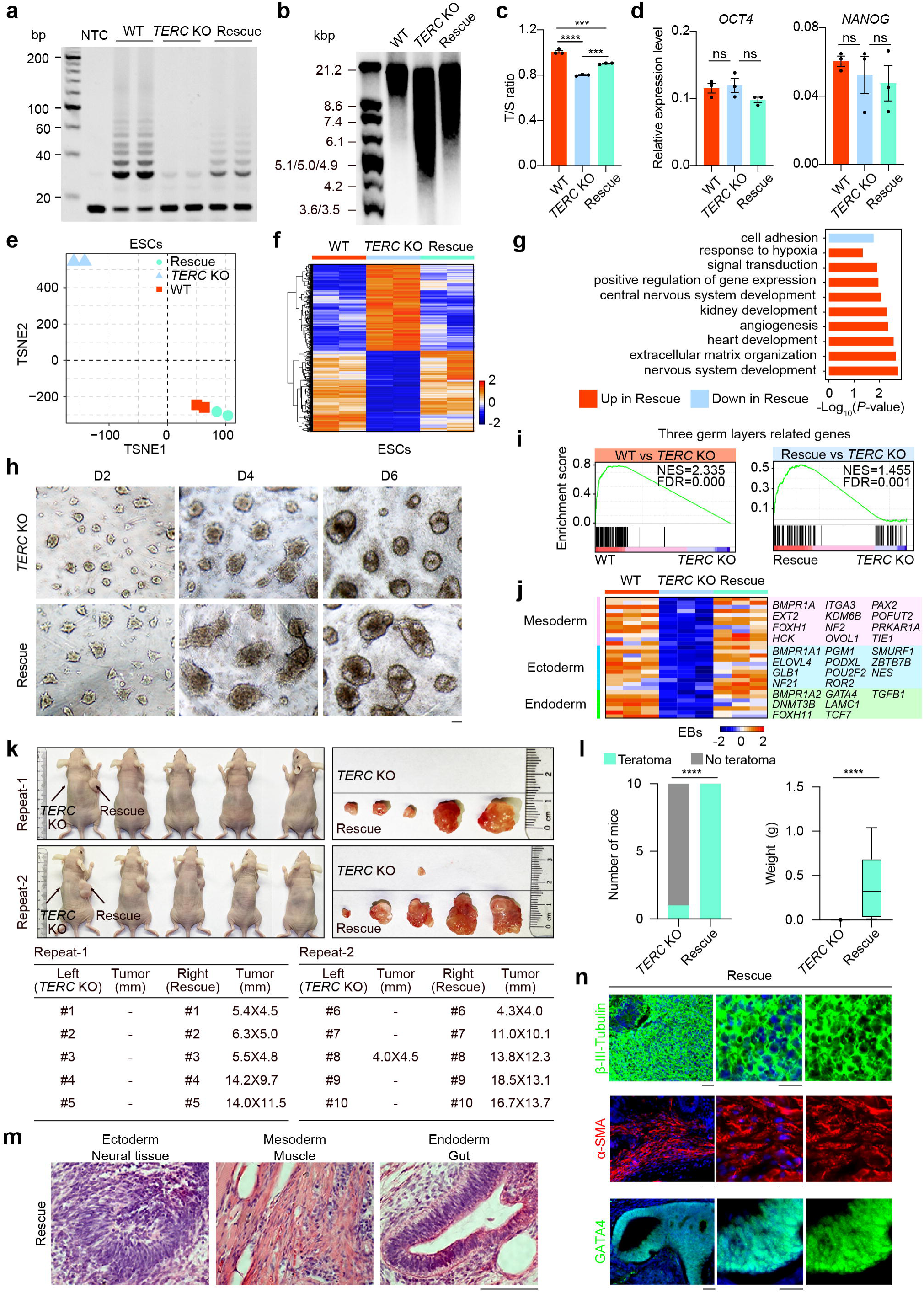
Reconstitution of *TERC* restores transcriptional programmes and lineage potential. **a,** TRAP assay for telomerase activity in WT, *TERC* KO and rescue hESCs at early passages. n=2 biological replicates. **b,** Absolute telomere length of WT, *TERC* KO and rescue hESCs, as measured via TRF. **c,** Relative telomere length of WT, *TERC* KO and rescue hESCs, as detected via the T/S ratio. The data are shown as the mean ± SEM; n=3 biological replicates. \*\*\**P* < 0.001, \*\*\*\**P* < 0.0001, ANOVA with multiple comparisons. **d,** qRT‒PCR analysis of the expression levels of *OCT4* and *NANOG* in WT, *TERC* KO and Rescue hESCs. The data are presented as the mean ± SEM; n=3 biological replicates. **e,** TSNE analysis of the transcriptomic profiles of WT, *TERC* KO and rescue hESCs. **f,** Heatmap depicting the expression profiles of rescued genes in WT, *TERC* KO and *TERC* rescue hESCs. **g,** GO enrichment of genes differentially expressed between *TERC* Rescue hESCs and *TERC* KO hESCs. **h,** Representative images of EBs derived from *TERC* KO and rescue EBs at days 2, 4, and 6 of *in vitro* differentiation. Scale bar, 100 μm. **i,** GSEA showing enrichment of germ layer-associated gene sets in WT (left) and *TERC* rescue (right) EBs. Red and blue represent genes whose expression is upregulated and downregulated, respectively, in WT/ *TERC* rescue EBs. **j,** Heatmap depicting the expression levels of genes associated with the three germ layers in WT, *TERC* KO and Rescue EBs. **k,** *In vivo* teratoma formation from *TERC* KO and *TERC* rescued hESCs in immunodeficient NU/NU mice. The arrows indicate the injection sites. n=10 biological replicates. Images of isolated teratomas 8 weeks after subcutaneous injection are shown. **l,** Quantification of teratoma formation in immunodeficient NU/NU mice 8 weeks after subcutaneous injection of *TERC* KO or rescue hESCs (left panel); the *P* value was calculated via the chi-square test. Box plot showing the weight of isolated teratomas (right panel); n=10 biological replicates; the *P* value was calculated via the Mann‒Whitney test. **m,** H&E staining of teratomas derived from *TERC* Rescue hESCs showing three germ layers, including neural tissue (ectoderm), muscle (mesoderm), and the gut (endoderm). Scale bar, 100 μm. **n,** Immunofluorescence staining for germ layer markers, including β-III-tubulin (ectoderm), α-SMA (mesoderm), and GATA4 (endoderm), from *TERC* rescue hESC-derived teratomas. Scale bar, 20 μm.

Given that complementing *TERC* in *TERC* KO hESCs effectively restored the expression of development-related genes and functions, we next sought to investigate whether *TERC* reintroduction could rescue the impaired differentiation capacity of *TERC* KO hESCs. First, we performed *in vitro* differentiation experiments and reported that restoring *TERC* could effectively restore the proliferation ability of EBs and that the morphology and size of EBs were similar to those of WT EBs (Fig. 3h and Fig. 2f). Transcriptome analysis of day 7 EBs revealed 1,374 upregulated and 1,583 downregulated genes in rescued *TERC* hESCs (Extended Data Fig. 4d), with approximately 40% of these DEGs showing rescued expression patterns (Extended Data Fig. 4e, f). Moreover, functional enrichment analysis revealed that re-introduction of *TERC* rescued multiple biological processes, particularly those related to development, including angiogenesis, nervous system development, and brain development (Extended Data Fig. 4g), which indicated that re-introduction of *TERC* for only 48 h reshaped the transcriptional landscape essential for proper *in vitro* differentiation. In addition, *TERC* KO resulted in the downregulation of key genes across the three germ layers, and conversely, *TERC* reintroduction reversed this effect on most genes (Fig. 3i, j). Most significantly, *in vivo* differentiation assays demonstrated that, unlike *TERC* KO, *TERC* re-introduction enabled proper lineage differentiation with three complete germ layer structures and appropriate expression of three germ layer-specific markers according to the results of a standard teratoma formation assay (Fig. 3k–n). Overall, the re-introduction of *TERC* in *TERC* KO hESCs reversed the impairment of differentiation potential, revealing the critical role of *TERC* involved in germ layer development.

### *TERC* orchestrates the transcriptional program of differentiation and development

To further investigate the potential mechanisms underlying the deficient differentiation capacity of *TERC* KO hESCs, we performed ChIRP-seq of *TERC* in hESCs as previously described^38^. Initially, a total of 9,945 *TERC*-specific binding regions were identified via unique mapping and strict screening (Fig. 4a). Further analysis revealed that many of these sites are located near transcription start sites (TSSs) and distal intergenic regions (Fig. 4b), and notably, approximately half of the regions targeted by *TERC* RNA are important regulatory elements containing promoters and enhancers (Extended Data Fig. 5a, b). These results provide solid evidence that *TERC* regulates gene transcription by targeting promoters and enhancers. *TERC*-bound sequences preferentially showed motif enrichment of key transcription factors, such as the Sp/KLF family (Fig. 4c), which are known to contain GC-rich motifs and regulate genes involved in differentiation and developmental processes^39^.

**Fig. 4.**
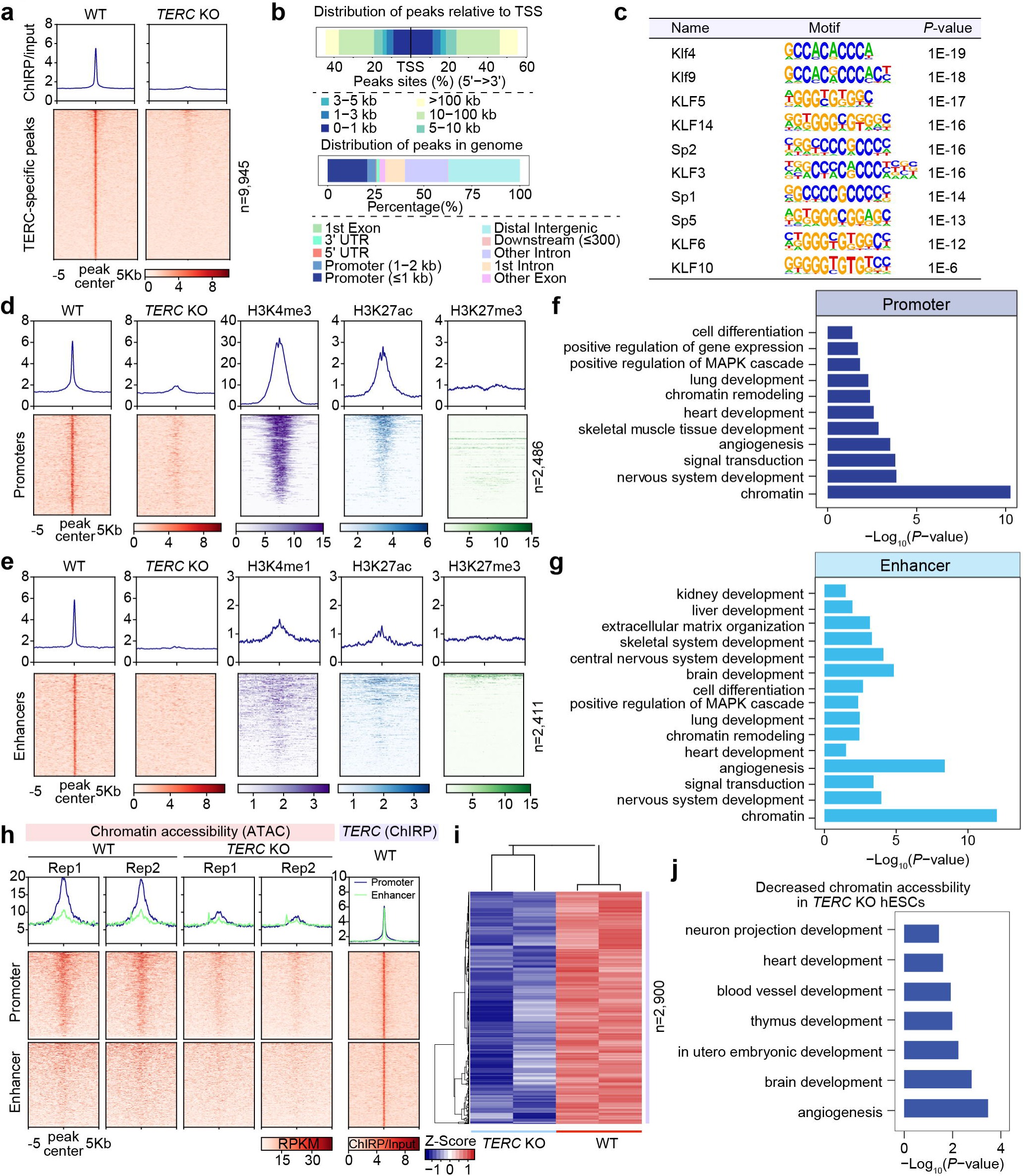
*TERC* occupies developmental regulatory elements associated with active chromatin states. **a,** Heatmap of ± 5 kb around *TERC* ChIRP-seq peaks (n=9,945) in WT vs. *TERC* KO hESCs. **b,** Bar plots showing the binding patterns of *TERC* peaks in the genome. **c,** Motif enrichment identified from *TERC* peaks. **d,** Heatmap showing enrichment of *TERC* and chromatin features at *TERC*-occupied promoters (n=2,486). **e,** Heatmap showing enrichment of *TERC* and chromatin features at *TERC*-occupied enhancers (n=2,411). **f,** Functional enrichment analyses for genes marked by *TERC* peaks distributed in promoters. These peaks were not enriched in *TERC* KO hESCs compared with those in WT hESCs. **g,** Functional enrichment analyses for genes marked by *TERC* peaks distributed in enhancers. **h,** Heatmap showing the chromatin accessibility of *TERC*-bound promoters (n= 2,486) and enhancers (n= 2,411) in WT and *TERC* KO hESCs. **i,** Heatmap of 2,900 ATAC-seq peaks significantly reduced in *TERC* KO hESCs. **j,** Functional enrichment analyses of genes marked by ATAC-seq peaks distributed in promoters and enhancers revealed significantly reduced accessibility in *TERC* KO hESCs compared with WT hESCs. Representative development-related terms from this analysis are shown.

Interestingly, *TERC*-bound regions were marked by the active histone modifications H3K4me3 and H3K27ac but lacked the repressive mark H3K27me3, indicating a transcriptionally permissive state (Fig. 4d, e). These *TERC*-bound sites were associated with genes involved in the MAPK, Wnt, and Hippo signalling pathways, as well as cancer-related pathways (Extended Data Fig. 5c,d). More importantly, genes targeted by *TERC*-bound promoters and enhancers were enriched in differentiation- and development-related terms, including nervous system development and angiogenesis (Fig. 4f, g and Extended Data Fig. 5e, f). Thus, *TERC* controls gene expression programs essential for differentiation and development through direct binding to promoters and enhancers.

### *TERC* promotes chromatin accessibility at regulatory regions of differentiation and developmental genes

Next, we asked whether *TERC* occupancy is associated with changes in chromatin accessibility, especially for genes related to development and differentiation. Biological replicates of ATAC-seq data from WT and *TERC* KO hESCs were highly reproducible, and *TERC* knockout resulted in a marked global reduction in chromatin accessibility (Extended Data Fig. 6a, d). Numerous ATAC-seq peaks were lost following *TERC* deletion, particularly at promoter and enhancer regions (Extended Data Fig. 6b), accompanied by reduced ATAC signal intensities (Extended Data Fig. 6c). Chromatin accessibility at TSSs was also decreased in *TERC* KO hESCs (Extended Data Fig. 6d), suggesting widespread transcriptional repression. Notably, *TERC*-specific binding sites exhibited reduced accessibility upon *TERC* knockout (Extended Data Fig. 6e), particularly at *TERC*-bound promoters, where the reduction was more pronounced than that at enhancers (Fig. 4h), revealing that *TERC* plays a pivotal role in maintaining an open chromatin state.

We performed a differential analysis of the ATAC-seq data and identified 2,900 ATAC-seq peaks with significantly reduced enrichment (Fig. 4i), which predominantly overlapped with promoters and enhancers (Extended Data Fig. 6f). Furthermore, these altered ATAC-seq peaks were enriched in active chromatin states (Extended Data Fig. 6g). Moreover, functional enrichment analysis revealed that these peak-marked genes were involved in developmental processes, such as brain and heart development, angiogenesis, and neuron development (Fig. 4j), and overlapped with regulators of the Wnt pathway (Extended Data Fig. 6h). Consistent with the ChIRP-seq results, motif analysis revealed enrichment of transcription factors such as NFY and the Sp/KLF family (Extended Data Fig. 6i), which are involved in the development of lineages, tissues and organs^39,40^. Taken together, these findings indicate that *TERC* is essential for maintaining the chromatin accessibility and transcriptional programs required for development.

### *TERC* is essential for H3K27ac deposition on differentiation- and development-related genes

Further analysis of regions with reduced chromatin accessibility revealed that these regions mainly coincided with H3K27ac-binding regions (Fig. 5a). Moreover, H3K27ac protein expression was significantly downregulated following *TERC* knockout (Extended Data Fig. 7a, b). Next, we performed chromatin immunoprecipitation sequencing (ChIP-seq) for H3K27ac in WT and *TERC* KO hESCs to determine changes in the genome-wide distribution of H3K27ac. H3K27ac occupancy in *TERC* KO hESCs resulted in extensive genome-wide loss (Fig. 5b and Extended Data Fig. 7c). Furthermore, identical motifs associated with three germ layers (*HOXA11*, *LHX1* and *LHX2*) in both ATAC-seq and H3K27ac ChIP-seq data indicated that the corresponding transcription factors may serve as critical bridges linking chromatin accessibility to epigenetic activation (H3K27ac) (Fig. 5c). *TERC* acts as a central driver of the entire differentiation process, and *TERC* knockout disrupts this functional bridge. Strikingly, H3K27ac occupancy was also significantly reduced at promoters and enhancers in *TERC* KO hESCs (Fig. 5d, e). Given the enrichment of *TERC* at promoters and enhancers, we examined H3K27ac occupancy in *TERC*-specific binding regions and found that these loci decreased similarly upon *TERC* knockout (Fig. 5f). Furthermore, chromatin accessibility at promoters and enhancers decreased after *TERC* was knocked out (Extended Data Fig. 6c), suggesting that *TERC* regulates chromatin accessibility by promoting H3K27ac deposition at developmental genes.

**Fig. 5.**
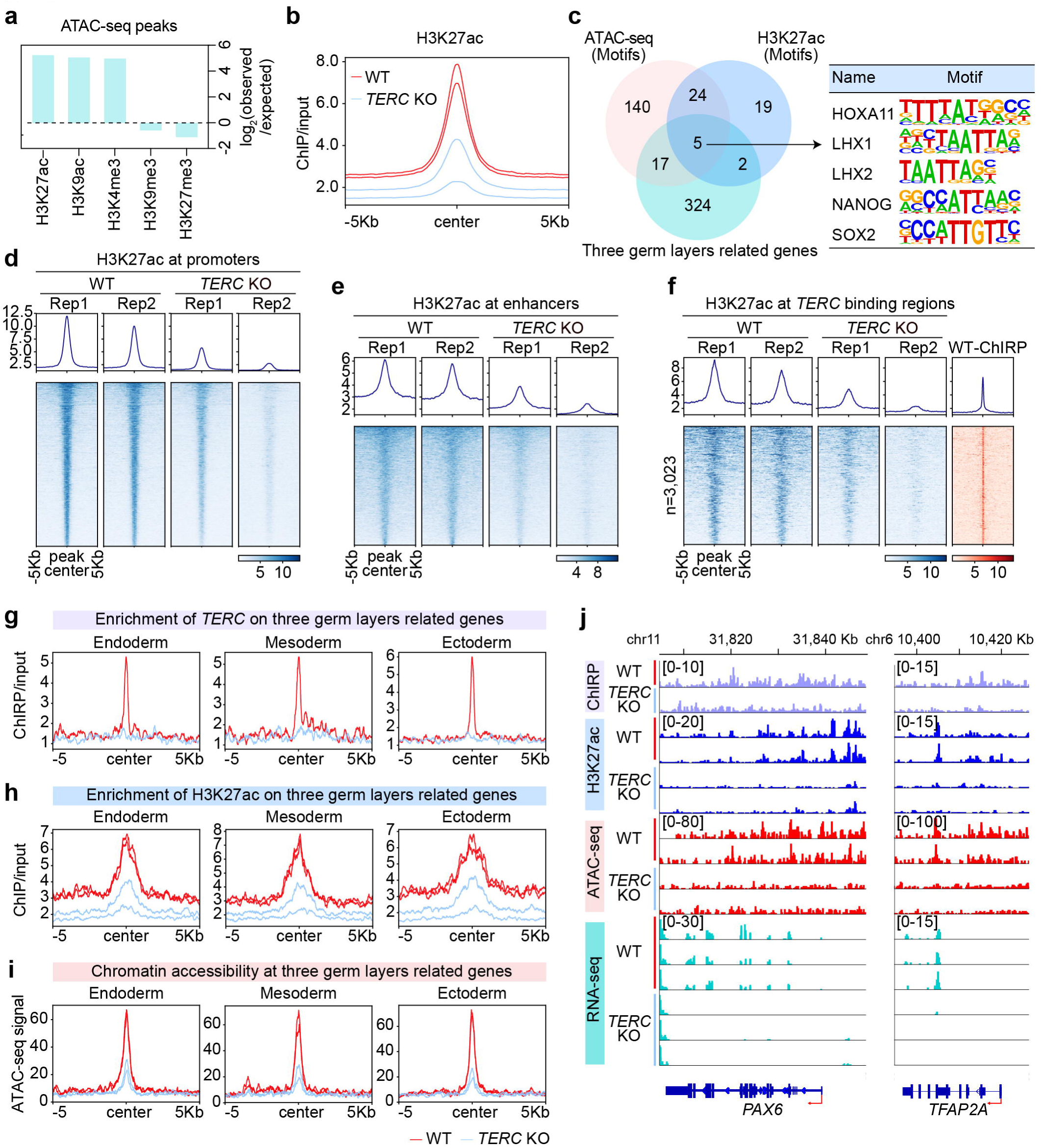
*TERC* promotes enhancer acetylation and chromatin accessibility at lineage-specifying genes. **a,** GAT analysis linking regions with reduced accessibility (n = 2,900) to histone modification-associated regions. The log2(observed/expected) represents the log2 of the fold enrichment value, given by the ratio of the observed count to the expected count. **b,** Average profile showing H3K27ac occupancy in WT and *TERC* KO hESCs at H3K27ac peaks. **c,** Motif analysis was performed on genomic regions defined by ATAC-seq or H3K27ac peaks that exhibited significantly reduced enrichment following *TERC* knockout, identifying key motifs involved in the three germ layers. **d,** Heatmap showing H3K27ac occupancy at promoters (TSS ± 2 kb) in WT and *TERC* KO hESCs. **e,** Heatmap showing H3K27ac occupancy at enhancers in WT and *TERC* KO hESCs. **f,** Heatmap showing H3K27ac occupancy at *TERC-*specific binding regions in WT and *TERC* KO hESCs. **g-i,** Average profile showing *TERC*, H3K27ac occupancy and ATAC-seq signals at three germ layer-related genes in WT and *TERC* KO hESCs. **j,** Genome browser representations of *TERC* ChIRP-seq, H3K27ac ChIP-seq, ATAC-seq and RNA-seq data at the *PAX6* and *TFAP2A* (ectoderm) loci upon *TERC* knockout. The red arrow indicates the direction of transcription.

Differential analysis revealed 5,040 H3K27ac peaks with reduced occupancy in *TERC* KO hESCs compared with 1,947 with increased occupancy (Extended Data Fig. 7d). Nearly half of the reduced peaks were located at promoters and enhancers (Extended Data Fig. 7e, f). Further functional enrichment analysis revealed that genes marked by these reduced H3K27ac peaks in promoters and enhancers were enriched for terms related to development, such as nervous system development, angiogenesis, brain development and heart development (Extended Data Fig. 7g). Moreover, *TERC* knockout led to a significant reduction in chromatin accessibility at these development-related genes (Extended Data Fig. 7h), suggesting that loss of H3K27ac leads to transcriptional silencing of development-related genes upon *TERC* knockout, thereby causing hESCs to exhibit impaired developmental potential.

Notably, our RNA-seq results confirmed that *TERC* knockout resulted in decreased expression of genes associated with lineage differentiation. *TERC* and H3K27ac binding to these genes was reduced upon *TERC* knockout, accompanied by decreased chromatin accessibility (Fig. 5g–i). For example, the expression levels of *PAX6* and *TFAP2A*, two genes that are markers of the neuroectoderm and nonneural ectoderm, respectively^41^, were downregulated in *TERC* KO hESCs, as *TERC* deficiency led to the loss of H3K27ac and decreased chromatin accessibility (Fig. 5j). In addition, the loss of H3K27ac caused by *TERC* knockout was involved in the regulation of key signalling pathways, including the MAPK and PI3K-Akt signalling pathways (Extended Data Fig. 7i–k). Together, these findings suggest that loss of *TERC* reduces chromatin accessibility and H3K27ac occupancy, resulting in failed lineage differentiation.

### *TERC* remodels TAD boundary integrity and chromatin domain architecture

To investigate whether *TERC* depletion alters higher-order genome organization, we performed *in situ* Hi-C in WT and *TERC* KO cells^42^. Hi-C libraries showed high mapping quality and reproducibility, with comparable mapping rates (∼80%), high cis-contact fractions, and expected strand orientation biases, indicating robust chromatin conformation capture quality (Extended Data Fig. 8a–c). Global compartment organization was first assessed using PC1 eigenvector decomposition at 100-kb resolution^43^. While overall A/B compartment proportions remained largely preserved between WT and *TERC* KO cells (Extended Data Fig. 8d, e), genome-wide compartment identity was extensively re-distributed upon *TERC* KO.

Comparisons of WT and *TERC* KO eigenvalues revealed substantial bidirectional compartment switching, including both A-to-B and B-to-A transitions (Fig. 6a). Quantitatively, ∼13–15% of the genomic bins underwent compartment switching, whereas ∼35% remained stably associated with either active (A) or inactive (B) compartments (Fig. 6b). Importantly, a saddle plot analysis^42,44^ demonstrated that long-range AA and BB compartmental interactions were globally maintained in *TERC* KO cells (Extended Data Fig. 8f), indicating that *TERC* KO remodels the distribution of chromatin states at the megabase scale while preserving the overall segregation principle of active and inactive chromatin domains. These data suggest that *TERC* KO does not induce a global collapse of chromatin compartmentalization but rather promotes the selective reassignment of compartment identity.

**Fig. 6.**
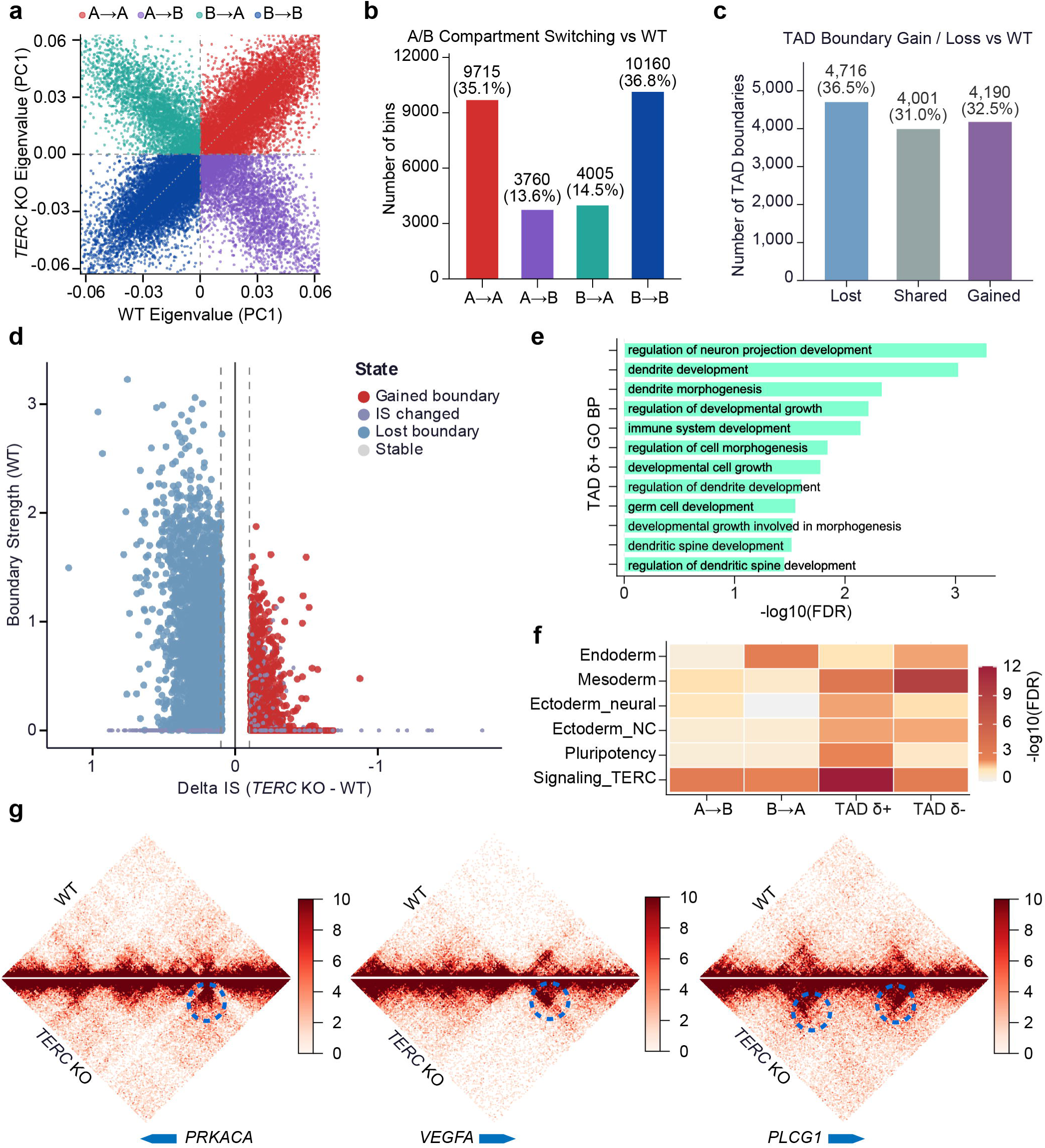
*TERC* contributes to higher-order chromatin organization in developmental domains. **a,** Scatter plot of first eigenvector (PC1) values comparing WT (x-axis) and *TERC* KO (y-axis) at 100 kb resolution. The genomic bins are colored according to compartment transition category: A→A (red), A→B (green), B→A (cyan), and B→B (blue). Dashed lines denote the A/B compartment boundary (PC1 = 0). The Pearson correlation coefficient is indicated. **b,** Bar chart showing the number of 100 kb genomic bins in each compartment transition category (A→A, 9,715 [35.1%]; A→B, 3,760 [13.6%]; B→A, 4,005 [14.5%]; B→B, 10,160 [36.8%]). In total, 28.1% of the genome underwent compartment switching upon *TERC* knockout. **c,** Bar chart showing the number of TAD boundaries at 50 kb resolution classified as lost (4,716; 36.5%), shared (4,001; 31.0%), or gained (4,190; 32.5%) upon *TERC* knockout. Boundaries were defined as local minima of the insulation score computed with a 250 kb sliding window. **d,** Scatter plot of the delta insulation score (ΔIS = ISKO − ISWT; x-axis) versus WT boundary strength (y-axis) for all 50 kb genomic bins. Points are colored by boundary state: gained boundary (red, ΔIS < −0.1 and boundary in KO), lost boundary (blue, ΔIS > +0.1 and boundary in WT), IS changed (purple, |ΔIS| > 0.1, other), and stable (gray, |ΔIS| ≤ 0.1). Dashed lines indicate the ΔIS threshold (± 0.1). **e,** Gene Ontology (GO) biological process enrichment analysis for genes overlapping genomic bins with lost/weakened TAD boundaries (TAD δ+, ΔIS > 0.1) in KOs. Bar lengths represent −log₁₀(FDR) (g:Profiler, gSCS correction). The most enriched terms were genes involved in neuronal morphogenesis and dendrite development. **f,** Heatmap of enrichment (−log₁₀ FDR, Fisher’s exact test) for curated developmental lineage gene sets across four classes of differential chromatin changes: A→B switching, B→A switching, TAD boundary weakening (TAD δ+), and TAD boundary strengthening (TAD δ−). Lineage categories: Endoderm, Mesoderm, Ectoderm_neural, Ectoderm_NC (neural crest), Pluripotency, and Signaling_*TERC*. **g,** Triangle Hi-C contact maps at three representative loci (*PRKACA*, *VEGFA*, and *PLCG1*) displaying altered chromatin interaction patterns upon *TERC* knockout. Upper triangle, WT; lower (inverted) triangle, KO. Balanced contact matrices at 50 kb resolution are shown on a linear scale (0–10, Red colormap). The genomic window spans ± 1.5 Mb flanking each gene body.

We next examined whether *TERC* KO affects local chromatin insulation and TAD organization^45,46^. Genome-wide TAD boundary analysis revealed extensive boundary remodelling following *TERC* KO, including both boundary loss and de novo boundary formation (Fig. 6c). Notably, lost boundaries represented the largest class of altered domains, accounting for 36.5% of all detected boundaries, whereas gained boundaries comprised 32.5% (Fig. 6c), suggesting that *TERC* KO preferentially destabilizes pre-existing insulated chromatin domains. To further quantify the insulation dynamics, we calculated the changes in the insulation score (IS) between WT and *TERC* KO cells^47^. Boundaries exhibiting increased IS values in *TERC* KO cells were classified as weakened or lost boundaries, whereas boundaries with reduced IS values were classified as strengthened or gained boundaries. Plotting ΔIS against WT boundary strength revealed that strong WT boundaries were disproportionately susceptible to insulation loss upon *TERC* KO (Fig. 6d). In contrast, newly gained boundaries generally emerged from regions with relatively weak preexisting insulation. These data suggest that *TERC* KO selectively disrupts highly insulated chromatin domains rather than inducing random architectural instability, which is consistent with the role of the targeted factor in maintaining higher-order chromatin insulation.

Genes linked to remodelled TADs were enriched for developmental and morphogenetic pathways, including those involved in neuron projection development, dendrite morphogenesis, developmental growth, and the regulation of cell morphogenesis (Fig. 6e). Lineage enrichment analysis further revealed strong associations between TAD remodelling events and developmental signalling programs, including ectodermal, mesodermal, and signalling-related transcriptional states (Fig. 6f). These observations suggest that *TERC* KO-mediated chromatin rewiring preferentially impacts regulatory regions associated with developmental plasticity and signalling responsiveness.

Inspection of representative genomic regions showed substantial local interaction remodelling in *TERC* KO cells, including altered contact enrichment surrounding the *PRKACA*, *VEGFA*, and *PLCG1* loci (Fig. 6g). These loci are associated with signalling, angiogenic, and growth-related pathways, suggesting that *TERC* KO preferentially rewires chromatin interactions at functionally responsive regulatory hubs. Consistent with these observations, loop analysis^42,48^ revealed a marked increase in the number of chromatin loops in *TERC* KO cells compared with WT cells (Extended Data Fig. 8g). Moreover, compared with those in WT cells, newly formed loops in *TERC* KO cells exhibited shorter genomic spans (Extended Data Fig. 8h), indicating enhanced local chromatin connectivity following the weakening of insulation.

Together, these findings support a model in which *TERC* KO redistributes chromatin topology across multiple hierarchical levels, weakening strongly insulated domains while promoting short-range regulatory interactions at developmental and signalling-associated genes.

## Discussion

Telomerase RNA acts as a template for telomere synthesis^1,2^. Here, we report that *TERC* is a global epigenetic regulator that is indispensable for the lineage differentiation of hESCs (Extended Data Fig. 9). The differentiation of hESCs is regulated by a dynamic interplay of epigenomic features, including chromatin accessibility, histone modifications, and transcription factor binding^24,25,49^. Our findings demonstrate that *TERC* is indispensable for the lineage differentiation of hESCs, primarily through its role in promoting active histone acetylation (H3K27ac) and chromatin accessibility.

*TERC* regulates differentiation by binding to the promoters and enhancers of developmental genes. Upon *TERC* depletion, hESCs exhibit a marked loss of H3K27ac at regulatory regions, resulting in collapsed chromatin accessibility and transcriptional repression. Our ChIRP-seq analysis revealed that *TERC* is associated with these regulatory elements of developmental genes, which are enriched in the motifs of key transcription factors (such as HOXA11, LHX1 and LHX2) that regulate lineage commitment. Consistently, a pioneering study revealed that *TERC* is associated with widespread genomic locations through the recognition of cytosine-rich consensus sequences^38^. The H2A–H2B heterodimer, as an RNA-binding factor for the CR4/5 domain of human *TERC*, also affects chromatin states^50,51^. *TERC* also regulates gene transcription by targeting the promoter regions of genes such as *LIN37*, *TPRG1L*, *TYROBP*, and *USP1*, thereby modulating inflammatory responses^17^. Additionally, *Terc* recruits RNA polymerase II to target myeloid gene promoters to increase myelopoiesis in zebrafish^52^. Our study highlights a model in which *TERC* binding is required to maintain a permissive chromatin state for the activation of developmental genes.

Human iPSCs can be generated from patients carrying heterozygous mutations in *TERC*^14,53^. *TERC*-mutated iPSCs exhibit defects in hematopoietic differentiation^53^. After sufficient passages, some iPSCs with fully activated OCT4 can upregulate the expression of *TERCs* and elongate telomeres^14^. Thus, reversing *TERC* defects or promoting *TERC* expression can aid in the treatment of *TERC* mutation-associated diseases. Consistently, engineered *TERC* RNA prevents senescence and resuces telomere length in iPSCs derived from patients carrying mutations in *TERC* and other telomere maintenance genes (*PARN* and *DKC1*)^54^. This mechanistic insight is corroborated by prior work demonstrating that PAPD5 inhibition elevates *TERC* levels and rescues hematopoietic defects in individuals with dyskeratosis congenita^55,56^. In line with these findings, we report that *TERC* re-introduction rescues both the expression of key developmental genes and their associated signalling pathways, demonstrating the essential role of *TERC* in maintaining an open chromatin state permissive for differentiation.

Our work refines the functionality of *TERC* as a global epigenetic regulator that is essential for stem cell fate by modulating the chromatin landscape, in addition to its canonical role as the RNA component of telomerase. However, the precise mechanism through which *TERC* regulates H3K27 localization directly or through intermediates remains to be investigated. While our Hi-C analyses revealed extensive TAD boundary remodelling upon *TERC* loss, whether *TERC* achieves chromatin localization through direct RNA‒DNA interactions, recognition of specific DNA sequence motifs, or bridging protein intermediaries—such as the H2A–H2B heterodimer known to bind the *TERC* CR4/5 domain^50,51^—remains important questions for future investigations. Additionally, transient *TERC* expression effectively rescues differentiation defects resulting from *TERC* deficiency. Slight telomere elongation is observed in *TERC* transient-expressing cells. We cannot formally exclude the contribution of slight telomere elongation observed in rescued cells. Future studies should employ *TERC* mutants to fully disentangle these effects. Furthermore, the functional conservation and divergence of *TERC* between species (e.g., mouse vs. human) and across stem cell types (embryonic vs. adult) remain to be systematically investigated. By revealing that a single noncoding RNA moonlights both telomere maintenance and epigenetic regulation, this work may have implications for deciphering the pleiotropic developmental phenotypes caused by *TERC* mutations.

## Methods

### Cell lines

The majority of the experiments were conducted using a female WA26 human embryonic stem (hES) cell line (WiCell, RRID: CVCL_E081) and a male H1 hES cell line (WiCell; RRID: CVCL_9771). hES cells were cultured under feeder-free conditions on growth factor-reduced (GFR) Matrigel-coated (1:100 dilution; Corning) plates and maintained in E8 medium (A1517001; Life Technologies) at 37°C with 5% CO2 under humidified conditions. hES cells were passaged every 3–4 days by incubation for 6 minutes with 0.5 mM EDTA in PBS at room temperature and replated in small clumps of cells in E8 medium. hES cells were routinely karyotyped, and their pluripotency and *in vitro* differentiation ability were tested by immunostaining and screening monthly for mycoplasma via PCR.

### Animal models

The animals used in this study were female Nu/Nu mice approximately 6 weeks of age and were purchased from Beijing Vital River Laboratory Animal Technology Co., Ltd. The mice were housed on a 12:12 hr light‒dark cycle with free access to food and water in individually ventilated specific pathogen-free (SPF) autoclaved cages at the College of Life Sciences. All animal housing, husbandry, and experimental procedures in this study were conducted in compliance with relevant guidelines and regulations, with approval from the Nankai University Animal Care and Use Committee.

### Plasmid construction

pSpCas9(BB)-2A-Puro (PX459, Addgene plasmid # 48139) and pSpCas9(BB)-2A-GFP (PX458, Addgene plasmid # 48138) were gifts from Feng Zhang. Guide RNAs were designed via the online design tool available at http://crispr.genome-engineering.org/. PX458/PX459 was digested with *Bbs*I and then gel purified. Two pairs of oligos, including targeting sequences, were annealed, and guide RNAs of *TERC* were cloned and inserted into *Bbs*I-digested PX458. *TERC* expression was rescued in *TERC* KO hESCs via transfection with pBabe-*TERC* or empty pBabe plasmids^17^, which were gifts from Prof. HaiYing Liu at Sun Yat-sen University. All relevant sgRNA sequences and primers are listed in Supplementary Table 1.

### Nucleofection

hESCs growing in Matrigel-coated dishes with E8 medium were detached with 0.5 mM EDTA. A total of 8 × 10^5^ cells were nucleofected with 10 μg of plasmids via Amaxa Nucleofector II (Lonza) and Human Stem Cell Nucleofector® Kits (Lonza) according to the manufacturer’s instructions. Nucleofected hESCs were plated back in a Matrigel-coated dish with E8 medium supplemented with 1 mM ROCK inhibitor. hESCs were subjected to puromycin selection 4 h after nucleofection and allowed to recover for 5 days.

To generate stable *TERC* KO hESCs, surviving (resistant) colonies were manually picked into new 24-well plates coated with Matrigel with E8 and then expanded for genotyping and sequencing. For the rescued *TERC* hESCs, colonies were directly expanded to detect *TERC* expression, and other experiments were performed after nucleofection and puromycin selection.

### Telomere Restriction Fragment Analysis

Telomere length was detected via telomere restriction fragment (TRF) analysis according to the commercial TeloTAGGG Telomere Length Assay (12209136001; Roche Life Science). Briefly, 1 μg of extracted genomic DNA was digested with a mixture of *Hinf I* and *Rsa I* enzymes at 37°C overnight. Next, the DNA fragments were separated on a 0.8% agarose gel at 6 V/cm in 0.5 × TBE buffer for 2.5 h. After denaturation and neutralization, the gels were transferred to a nylon membrane (RPN2020B; GE Healthcare) at 25°C for 24 h. The membrane was hybridized with digoxigenin (DIG) Easy Hyb Granules containing a telomere probe at 42°C overnight and then incubated with anti-DIG-alkaline phosphatase for 4 h. The telomere signal was detected via chemiluminescence after a substrate solution was added to the membrane.

### Telomere quantitative fluorescence *in situ* hybridization (Q-FISH)

Telomere length was detected via Q-FISH as described previously^22^. The cells were incubated with 0.3 μg/mL nocodazole for 4 h to enable metaphase of the cells. Metaphase-enriched cells were subjected to hypotonic treatment with 0.075 M KCl solution, fixed with methanol:glacial acetic acid (3:1) and spread onto clean slides. Telomeres were denatured at 80°C for 3 min and hybridized with a TelC-Cy3 probe (F1002; Panagene) at 0.5 μg/mL. Chromosomes were counterstained with 0.5 μg/mL DAPI. The fluorescence was detected and imaged via an Axio-Imager Z2 fluorescence microscope (Zeiss).

### Telomere measurement by the T/S ratio

The average telomere length was measured from total genomic DNA via real-time PCR^57^. PCR was performed via an iCycler MyiQ2 detection system (Bio-Rad) with primers targeting telomeres and the reference gene (human 36B4 single-copy gene) under previously described conditions (Table S2)^58^. For each PCR, a standard curve was generated via serial dilutions of known amounts of DNA. The telomere signal was normalized to the signal from the single-copy gene to generate a T/S ratio indicative of relative telomere length. Equal amounts of DNA (20 ng) were used for each reaction.

### Telomerase activity by TRAP assay

Telomerase activity was determined via the TRAP method according to the manufacturer’s instructions via a TeloChaser Telomerase assay kit (T0001; MD Biotechnology). Approximately 1 × 10^4^ cells from each sample were lysed, and lysis buffer served as a negative control. The PCR products of the cell lysates were separated by nondenaturing TBE-based 10% polyacrylamide gel electrophoresis and visualized via ethidium bromide staining.

### Western blot

The cells were washed twice in PBS, lysed in cell lysis buffer on ice for 30 min and then sonicated for 1 min at an amplitude of 60 at 2 s intervals. After centrifugation at 10,000 × g for 10 min at 4°C, the supernatant was transferred into new tubes. The protein concentration of each sample was measured via bicinchoninic acid, and the protein samples were boiled in SDS sample buffer at 100°C for 10 min. The protein of each cell extract was resolved via 10% Acr-Bis SDS‒PAGE and transferred to polyvinylidene difluoride membranes (Millipore). The membrane was blocked with 5% skim milk in TBST at room temperature for 2 h and then incubated with primary antibodies overnight at 4°C. β-actin served as a loading control. The immunoreactive bands were then probed for 2 h at RT with the appropriate horseradish peroxidase (HRP)-conjugated secondary antibodies. The protein bands were detected with a chemiluminescent HRP substrate (WBKLS0500, Millipore). The following antibodies were used for Western blotting: anti-OCT4 (Santa Cruz; sc-5279; 1:1000), anti-NANOG (Abcam; ab80892; 1:1000) and anti-β-actin (ABclonal; AC026; 1:50000).

### Immunofluorescence microscopy

The cells were subsequently washed twice with PBS, fixed with fresh 3.7% paraformaldehyde for 30 min at 4°C, permeabilized with 0.1% Triton X-100 in blocking buffer (3% goat serum plus 0.1% BSA in PBS) for 20 min at room temperature (RT), incubated with blocking buffer for 1 h at RT, and stained with primary antibodies overnight at 4°C. The cells were subsequently incubated with fluorescence-labelled secondary antibodies for 2 h at RT. Hoechst 33342 was used to stain the nuclear DNA. The fluorescence was detected and imaged via an Axio-Imager Z2 fluorescence microscope (Zeiss). The following antibodies were used for these experiments: anti-alpha smooth muscle actin (Abcam; ab5694; 1:200), anti-β-ΙΙΙ-tubulin (Abcam; ab78078; 1:200), anti-alpha 1 Fetoprotein (Abcam; ab213328; 1:200), anti-GATA4 (Santa Cruz; sc-25310; 1:200), donkey anti-rabbit IgG Alexa Fluor 594 (Thermo Scientific; A-21207; 1:200) and donkey anti-mouse IgG Alexa Fluor 488 (Thermo Scientific; A-21202; 1:200).

### RT‒qPCR

Total RNA was extracted from cells via an RNeasy RNA Mini Kit (74104; QIAGEN) according to the manufacturer’s instructions. Reverse transcription was performed on purified total RNA to generate cDNA via M-MLV reverse transcriptase (Invitrogen) and random hexamer primers (*TERC* RNAs) or Oligo (dT)18 primers according to the manufacturer’s instructions. qPCR was performed with FastStart Universal SYBR Green Master Mix (4913914001; Roche) on an iCycler MyiQ2 detection system (Bio-Rad). Each sample was set up in duplicate and normalized to the expression of GAPDH. qPCR data were analysed via the ΔΔCt method. qPCR primers were confirmed for their specificity via dissociation curves and are listed in Supplementary Table 2.

### Embryoid body formation

Embryoid body formation was carried out following established protocols^59^. After the cells were washed with PBS, they were dissociated with EDTA solution, and then a 1-mL pipette tip was used to spray the colonies with 1 mL of E8 medium to detach them from the plate. The mixture was triturated via a 1-mL pipette tip until the solution became cloudy with single cells, after which the cells were counted. The cells were centrifuged at 270 × g for 5 min, and the cells were first resuspended in 1 mL of low-bFGF hESC medium supplemented with ROCK inhibitor (1:100, final concentration 50 μM). The mixture was pipetted up and down to ensure a single-cell suspension. Next, an additional volume of low-bFGF hESC medium supplemented with ROCK inhibitor was added to obtain 9,000 live cells per 150 μL. A 150 μL suspension was dispensed into each well of a low-attachment 96-well U-bottom plate, followed by incubation at 37°C under 5% CO₂ for continued culture.

### Teratoma formation analysis

WT hESCs, *TERC* KO hESCs, and Rescued *TERC* hESCs were well maintained on Matrigel-coated 6-well plates. The hESCs were dissociated into single cells and resuspended in 50% Matrigel diluted in DMEM/F12 (Thermo Scientific). Next, 1 × 10^6^ hESCs were subcutaneously injected into immunodeficient NU/NU mice at approximately 6 weeks of age. Eight weeks later, the mice were humanely sacrificed, and the teratomas were isolated and fixed in 4% paraformaldehyde. Teratomas were stained with hematoxylin and eosin (H&E) or used for immunofluorescence. The experiments involving animal research for teratoma formation were reviewed and approved by the Nankai University Animal Care and Use Committee (NO. 2023-SYDWLL-000639).

### Hematoxylin and eosin (HE) staining

After deparaffinization in xylene (2 × 5 min) and rehydration through an ethanol series (100%, 85%, 70%; 5 min each), the sections were stained with hematoxylin (4 min), rinsed in ddH₂O (5 min), differentiated in 1.5% HCl-75% ethanol (4 sec), and washed in ddH₂O (5 min). Following PBS rinsing (2 min) and eosin counterstaining (20 sec), the sections were dehydrated through graded ethanol (70%, 85%, 95%, 100%), cleared in xylene, and mounted with neutral resin medium. All steps were conducted at room temperature.

### Fluorescence microscopy of the teratoma sections

Briefly, after being deparaffinized, rehydrated, and washed in 0.01 M PBS (pH 7.2–7.4), the sections were incubated with 3% H2O2 for 10 min at room temperature to block endogenous peroxidase activity, subjected to high-pressure antigen recovery sequentially in 0.01 M citrate buffer (pH 6.0) for 3 min, and permeabilized with 0.1% Triton X-100 for 1 h. Next, the sections were incubated with blocking solution (3% BSA in PBS) for 2 h at room temperature and then incubated with diluted primary antibodies overnight at 4°C. Blocking solution without the primary antibody served as a negative control. After being washed with PBS three times (15 min each), the sections were incubated with the appropriate fluorescence-conjugated secondary antibodies for 2 hours at room temperature. The sections were subsequently washed three times in PBS, counterstained with Hoechst 33342, and placed in Vectashield mounting medium (H-1000-10; VectorLabs). The fluorescence was detected and imaged via an Axio-Imager Z2 fluorescence microscope (Zeiss).

### RNA-seq library preparation and sequencing

Total RNA was extracted from cells via an RNeasy Mini Kit (QIAGEN) according to the manufacturer’s instructions. Briefly, mRNA was purified from total RNA via poly-T oligo-attached magnetic beads. Fragmentation was carried out using divalent cations under elevated temperature in First Strand Synthesis Reaction Buffer. First-strand cDNA was synthesized via random hexamer primers and M-MuLV Reverse Transcriptase (RNase H-). Second-strand cDNA synthesis was subsequently performed via DNA polymerase I and RNase H. The remaining overhangs were converted into blunt ends via exonuclease/polymerase activities. After adenylation of the 3’ ends of the DNA fragments, adaptors with hairpin loop structures were ligated to prepare for hybridization. The library fragments were subsequently purified with the AMPure XP system. PCR was subsequently performed with Phusion High-Fidelity DNA polymerase, universal PCR primers and Index Primer. Finally, the PCR products were purified. The final indexed libraries were pooled and sequenced on an Illumina NovaSeq platform with a 150-bp paired-end read length.

### ATAC-seq library preparation and sequencing

The ATAC-seq libraries of hESCs were prepared as previously described with minor modifications^60,61^. For each sample, 500 cells were counted, washed with cold PBS, and subsequently lysed in cold lysis buffer (10 mM Tris-HCl (pH 7.4), 10 mM NaCl, 3 mM MgCl2 and 0.2% NP-40) on ice to prepare the nuclei. Immediately after lysis, the nuclei were spun at 600 × g for 5 min to remove the supernatant. The nuclei were then incubated with Tn5 transposome and tagmentation buffer at 37°C for 30 min (Vazyme Biotech). After tagmentation, stop buffer was added directly to the reaction mixture to stop tagmentation. PCR was performed to amplify the library for 14 cycles via the following PCR conditions: 72°C for 3 min; 98°C for 30 s; and thermocycling at 98°C for 15 s, 60°C for 30 s and 72°C for 3 min, followed by 72°C for 5 min. After PCR, the libraries were purified with AMPure beads (Beckman), and the beads were subsequently eluted in 25 µL of ddH2O. The final indexed libraries were pooled and sequenced on an Illumina HiSeq 4000 platform with a 150-bp paired-end read length.

### ChIRP-seq

ChIRP-seq was carried out as previously described^38^. Briefly, 2 × 10^7^ hESCs were crosslinked with 3% formaldehyde for 30 min at room temperature and then quenched with 0.125 mM glycine for 5 min. After being rinsed with chilled PBS, the cells were snap frozen in liquid nitrogen and stored at −80°C. The cell pellets were resuspended in cell lysis buffer containing 50 mM Tris–HCl (pH 7.0), 10 mM EDTA, and 1% SDS; 1 mM PMSF (Sigma‒Aldrich), protease inhibitors (Roche) and Superase-in (Thermo Scientific) were freshly added; and the samples were subsequently sonicated until most of the DNA was smaller than 500 bp in size. The lysate was centrifuged to remove insoluble material, and the supernatant was transferred to fresh tubes. Two inputs were saved for RNA enrichment analysis via RT–qPCR and DNA samples. Hybridization was performed in a 2× volume of freshly prepared hybridization buffer (750 mM NaCl, 1% SDS, 50 mM Tris–HCl [pH 7.0], 1 mM EDTA, 15% formamide, 1 mM PMSF, protease inhibitors, and Superase-in) supplemented with 1 µL of 100 µM *TERC* odd probes at 37°C with end‒end rotation for 16 h. Streptavidin C1 beads (Invitrogen) were then added, followed by an additional incubation of 30 min at 37°C with shaking. The beads were subsequently washed five times with wash buffer (2 × SSC, 0.5% SDS and 1 mM PMSF) at 37°C for 5 min. RNA and DNA were isolated separately and subjected to qPCR analysis or high-throughput sequencing.

### ChIP-seq library preparation and sequencing

ChIP-seq was performed as previously described^62,63^. The collected hESCs were crosslinked with 1% formaldehyde for 10 min, quenched with glycine, and lysed. Chromatin was sonicated to generate 100–500 bp fragments and immunoprecipitated overnight at 4°C with H3K27ac antibody (Abcam; ab177178) and Dynabeads M280 (Life Technologies). After sequential washes, the complexes were eluted from the beads by heating for 15 min at 65°C and then reverse crosslinked overnight at 65°C. DNA was treated with RNase A and proteinase K and extracted with phenol:chloroform:isoamyl alcohol (25:24:1, pH > 7.8). The purified DNA was subjected to the NEBNext® Ultra^TM^ II DNA Library Prep Kit (NEB, USA, E7645L) for ChIP-seq library preparation. The final DNA library was analysed for size distribution via an Agilent 5400 system, and the qualified libraries were pooled and sequenced on an Illumina NovaSeq platform with the PE150 strategy at Novogene Bioinformatics Technology Co., Ltd. (Beijing, China).

### *In situ* Hi-C library preparation

Hi-C libraries were prepared largely as described previously^64^. Briefly, cells were crosslinked with 2% formaldehyde and lysed to isolate nuclei. Chromatin was then digested overnight with DpnII. The resulting restriction fragment overhangs were filled in with a mix of biotin-14-dATP, and the DNA ends were marked. After inactivating the enzymes, the blunt-ended fragments were ligated *in situ* under dilute conditions using T4 DNA ligase to favour ligation between crosslinked fragments. Following crosslink reversal and DNA purification, the DNA was sheared to 200–400 bp fragments using a Bioruptor. Biotin was removed from unligated, nonproximal ends, and the DNA containing biotin-labelled ligation junctions was specifically captured using streptavidin magnetic beads. Libraries were then constructed on-bead using the NEBNext Ultra DNA Library Prep Kit. The final libraries were size-selected for a range of 200–600 bp and sequenced on an Illumina HiSeq platform.

### RNA-seq data processing

The raw RNA-seq data with low-quality read pairs and adapters were trimmed via TrimGalore to obtain clean reads. Next, the trimmed clean reads were aligned to the human reference genome hg19 via HISAT2 with the default settings^65^. The uniquely mapped reads unambiguously attributed to each gene were counted with featureCounts^66^. Differential expression analysis was conducted via the DESeq2 package in R^67^, and adjusted *P* values were computed in DESeq2 via the Wald test and adjusted for multiple testing via the procedure of Benjamini and Hochberg^68^. Differentially expressed genes (DEGs) were identified via the following criteria: for WT versus *TERC* KO comparisons, fold change > 1.5 and adjusted *P* value < 0.05; and for *TERC* KO versus Rescued *TERC* comparisons, fold change > 1.2 and adjusted *P* value < 0.05. Gene Ontology (GO) and Kyoto Encyclopedia of Genes and Genomes (KEGG) analyses of the DEGs were performed via DAVID, and only terms whose *P* value was < 0.05 were considered significantly enriched^69^. GSEA was conducted to identify enrichment, and only gene sets with an FDR < 0.05 were considered significantly enriched^70^.

### ATAC-seq and ChIP-seq data processing

For ATAC-seq data, adapter sequences were trimmed, and low-quality or low-complexity read pairs were filtered from the raw data via TrimGalore. Afterward, the qualified read pairs were aligned to the human reference genome hg19 via Bowtie2 with default parameters^71^, which retained only the alignments with the best mapping quality. The duplicate reads were removed via Sambamba^72^. ATAC peaks were called via MACS2 with the parameters “-q 0.01 -g hs -f BAMPE’’^63,73^. To minimize background noise, the IDR package was employed to identify highly reproducible peaks^74^. Differential binding analysis of ATAC-seq or ChIP-seq peaks was conducted with the DiffBind package. DeepTools was used to visualize histone modification signals or chromatin accessibility around peaks or genes^75^.

### ChIRP-seq data processing

The analysis pipeline for ChIRP-seq was previously published^38,76^. Briefly, the ChIRP-seq data were aligned to the human reference genome hg19 via Bowtie2^71^. The peaks of each sample were called via MACS2 against its corresponding input^73^. DeepTools was used to visualize *TERC* binding signals around peaks or genes^75^.

### Hi-C data analysis

Hi-C sequencing data were processed using a custom Snakemake-based pipeline^77^. The raw reads were quality controlled (FastQC/MultiQC), trimmed (Trim Galore), and aligned to the GRCh38 reference genome using BWA-MEM^78^. The resulting alignments were processed with pairtools to parse valid contacts (MAPQ ≥ 30), remove PCR duplicates, and select high-quality pairs^79^. After passing library quality control thresholds for metrics such as cis-contact ratios, the data were used to construct contact matrices at 15, 50, and 100 kb resolutions, which were subsequently ICE-normalized using a cooler^80,81^. Changes in chromatin architecture between WT and KO mice were assessed at multiple levels. A/B compartments were identified at 100 kb resolution using the eigenvector (PC1) method in cooltools^82^, with compartment switching and strength changes evaluated^43^. Topologically associating domain (TAD) boundaries were called at 50 kb resolution via the insulation score method^47^, and differential boundaries were classified as gained, lost, or weakened/strengthened on the basis of changes in the insulation score (ΔIS). Chromatin loops were identified at 15 kb resolution using Mustache^83^, after which differential analysis and aggregate peak analysis (APA) were performed to assess loop strength^42^. To elucidate the biological significance of these structural alterations, Gene Ontology (GO) and developmental lineage enrichment analyses were performed on genes located within regions that exhibited significant changes^84,85^, such as compartment switching or altered TAD insulation. Key interaction matrices were visualized as triangular heatmaps for direct comparison.

### Annotation and enrichment analysis of genomic regions

The ‘‘annotatePeak’’ function in the R package ChIPseeker was utilized for peak annotation to characterize their genomic features^86^. In accordance with a previously published study^87^, the promoters were defined as the ± 2 kb regions flanking transcription start sites (TSSs), and the active promoters were those associated with H3K4me3 but not H3K27me3. Enhancers were subsequently identified as H3K4me1 peaks, excluding those overlapping with promoter regions, and active enhancers were defined as distal H3K4me1 peak regions with H3K27ac but not H3K27me3. Motif enrichment analysis was performed on selected genomic regions via Homer’s ‘findMotifsGenome’^88^.

Functional enrichment analysis of genes associated with or proximal to the identified peaks was performed via DAVID, with only terms showing statistically significant enrichment (*P* < 0.05) being considered. The fold enrichment, represented as log2(observed/expected), was computed via the Genomic Association Test (GAT) software^89^.

### Statistical analysis

Statistical analyses were performed via an unpaired two-tailed Student’s *t* test in PRISM software (GraphPad 8 Software) to compare the differences between the treatment and control groups, with the assumption of equal variance. The Mann‒Whitney test was used to determine the significance of differences between data without a normal distribution. One-way or two-way ANOVA with Tukey’s test was used for multiple comparisons. The chi-square test was used to test differences between two groups for categorical variables. *, **, *** and **** indicate *P* < 0.05, *P* < 0.01, *P* < 0.001 and *P* < 0.0001, respectively. NS indicates not significant.

## Data availability

All sequencing data reported in this paper have been deposited in the Genome Sequence Archive (GSA) of the National Genomics Data Center, China National Center for Bioinformation/Beijing Institute of Genomics, Chinese Academy of Sciences, under accession number HRA013045 (BioProject accession: PRJCA045388), which can be accessed at URL: https://ngdc.cncb.ac.cn/gsa-human/s/eDS56rN3. The raw data can be requested via the GSA-Human System and can be authorized for download by the Data Access Committee for research and noncommercial use only. This paper does not report the original code. Any additional information required to reanalyze the data reported in this paper is available from the corresponding author upon reasonable request.

## Author contributions

J.L. and H.W. designed the experiments, conducted the major experiments, analysed the data, and prepared the manuscript. P.S. and M.G. performed the Hi-C experiments and data analysis, and C.L. conducted the teratoma formation assay-related experiments with assistance from N.L.; P.S., M.G., and C.L. all revised the manuscript. H.L. provided the pBabe-*TERC* plasmid to generate rescued *TERC* hESCs. G.F., Y.Y., Z.C., X.Y., and Z.J. conducted and discussed the immunofluorescence staining experiments of teratoma sections. G.Y., Z.M.Z. and J.T.L. assisted with portions of the bioinformatics analysis. L.L. and H.W. conceived the project, designed the experiments, and revised the manuscript.

## Additional information

Extended data Fig. 1–9

Supplementary Table 1. Primers for the CRISPR/Cas9 experiments

Supplementary Table 2. Primers for the T/S ratio and RT‒qPCR

## Extended data

**Extended Data Fig. 1.**
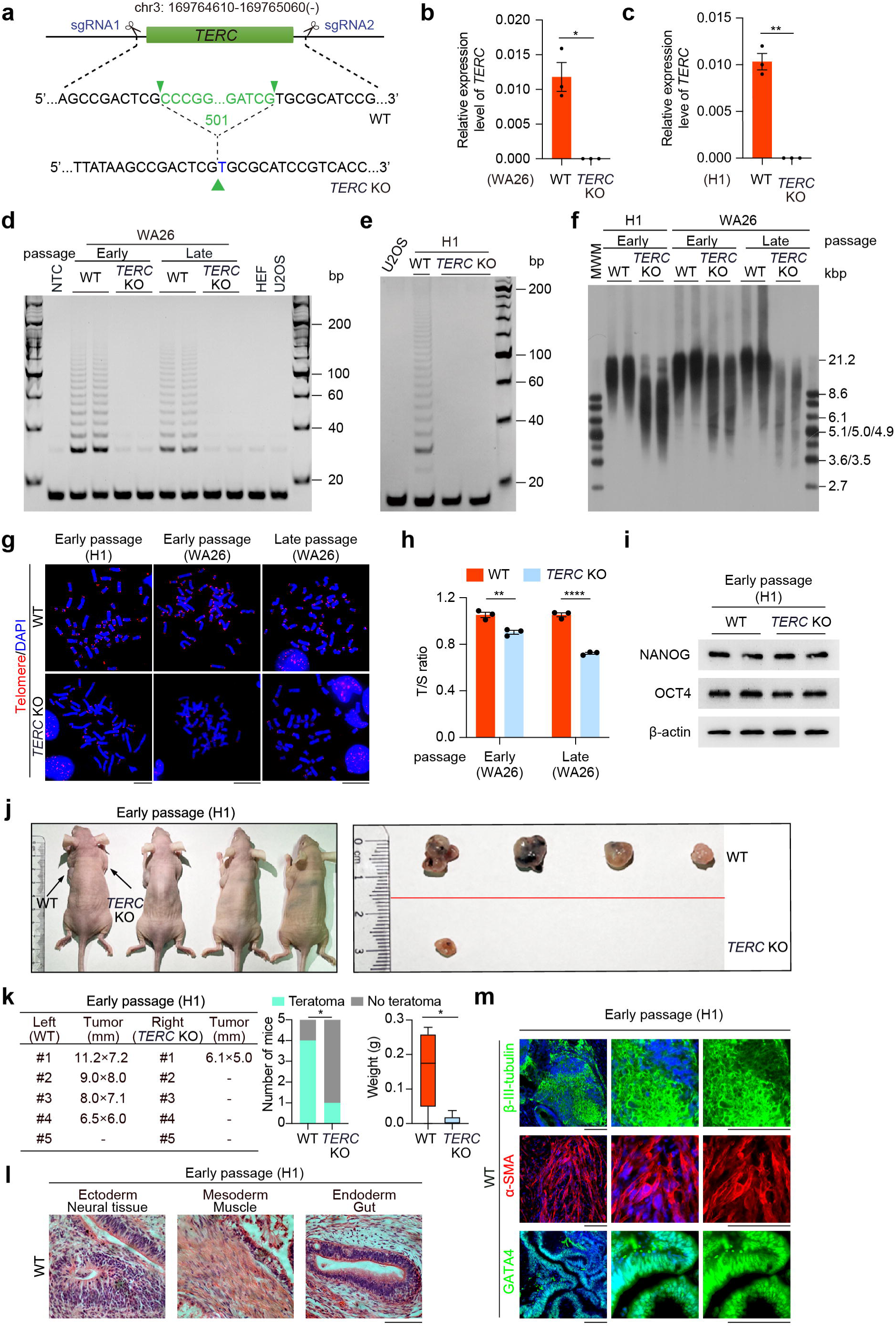
Knockout of *TERC* impairs the differentiation potential of hESCs. **a,** Schematic representation of CRISPR-Cas9 mediated knockout of *TERC* in hESCs. Successful editing was confirmed by Sanger sequencing. **b, c**, qRT‒PCR analysis of the expression levels of *TERC* RNA in *TERC* KO hESCs (WA26 and H1) compared with those in WT hESCs. n=3 biological replicates, means ± SEMs, and *P* values were calculated via two-tailed unpaired t tests. **d,** Telomerase activity was assessed by a TRAP (telomeric repeat amplification protocol) assay in *TERC* KO hESCs (WA26; early passage: P8 and late passage: P20) compared with WT hESCs. n=2 biological replicates. HEF and U2OS cells served as telomerase negative controls. **e,** TRAP assay for telomerase activity in *TERC* KO hESCs (H1; early passage: P8) compared with WT hESCs. U2OS cells served as telomerase negative controls. **f,** Telomere restriction fragment (TRF) analysis comparing telomere lengths in *TERC* KO and WT hESCs (H1 and WA26; early passage: P8 and late passage: P20). n=2 biological replicates. **g,** Representative telomere FISH images of *TERC* KO and WT hESCs (H1 and WA26; early and late passages). Blue, DAPI-stained chromosomes. Red, telomeres. Scale bar, 10 μm. **h,** The T/S ratio was used to measure the relative telomere length of *TERC* KO and WT hESCs (WA26; early and late passages). n=3 biological replicates, means ± SEMs, and *P* values were calculated via two-tailed unpaired t tests. **i,** Western blot analysis of pluripotent factors (OCT4 and NANOG) in *TERC* KO and WT hESCs (H1; early passages). n=2 biological replicates. **j-k,** *In vivo* differentiation of *TERC* KO and WT hESCs (H1; early passage) by teratoma assays in immunodeficient Nu/Nu mice (**j**). n=5 biological replicates; the chi-square test was used to test differences between two groups for categorical variables. The size and weight of the teratomas 8 weeks after subcutaneous injection into the left or right of NU/NU mice with the indicated hESCs are shown (**k**). The weights of the teratomas are shown in box plots, and the *P* value was calculated via the Mann‒Whitney test. **l,** Hematoxylin and eosin staining of teratomas from WT hESCs (H1; early passage). Teratomas consist of representative derivatives of three germ layers, including neural tissue (ectoderm), muscle (mesoderm), and the gut (endoderm). Scale bar = 100 μm. **m,** Immunofluorescence staining of markers representing three germ layers from WT hESC-derived teratomas (H1; early passage), including β-III-tubulin (ectoderm), α-SMA (mesoderm), and GATA4 (endoderm). Scale bar = 100 μm.

**Extended Data Fig. 2.**
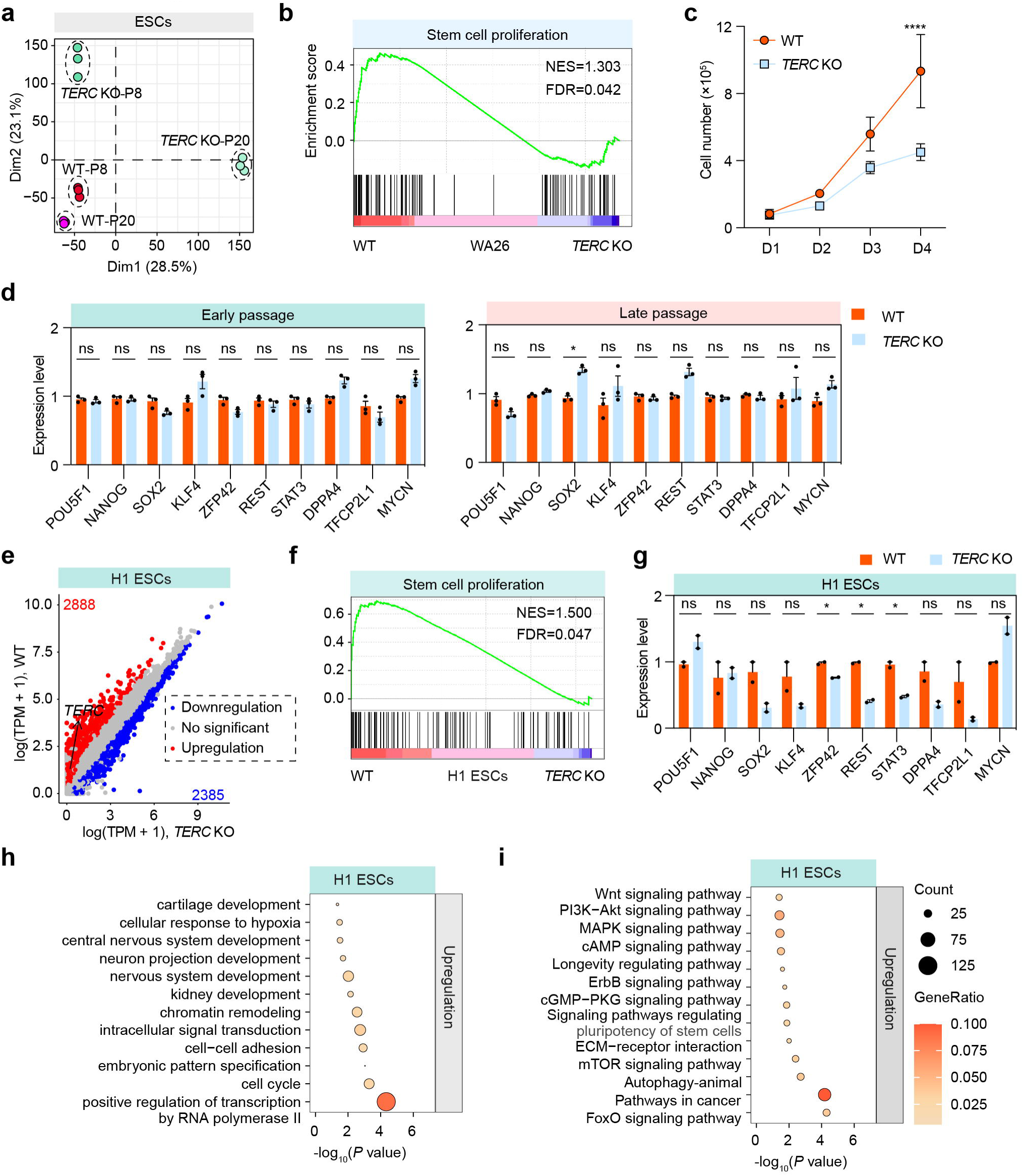
*TERC* deficiency impairs development-related signalling pathways and biological processes. **a,** Principal component analysis (PCA) of RNA-seq data from WT and *TERC* KO hESCs (WA26; early and late passages). **b,** GSEA indicating that upregulated genes in WT hESCs (WA26) were highly enriched in the gene set associated with stem cell proliferation. **c,** Proliferation curves of WT and *TERC* KO hESCs (WA26; early passage) over 4 days. Two-way ANOVA with multiple comparisons, *P* < 0.0001. **d,** RNA-seq analyses were performed to determine the expression levels of pluripotency markers in WT and *TERC* KO hESCs (WA26; early and late passages). **e,** Scatterplot showing differential gene expression levels between WT and *TERC* KO hESCs (H1; early passage). The red and blue dots represent genes whose expression was upregulated and downregulated, respectively, in WT hESCs compared with *TERC* KO hESCs. **f,** GSEA indicating that upregulated genes in WT hESCs (H1) were highly enriched in the gene set associated with stem cell proliferation. **g,** RNA-seq analyses were performed to determine the expression levels of pluripotency markers after *TERC* was knocked out in hESCs (H1; early passage). **h, i,** GO and KEGG analyses of upregulated genes in WT hESCs (H1; early passage).

**Extended Data Fig. 3.**
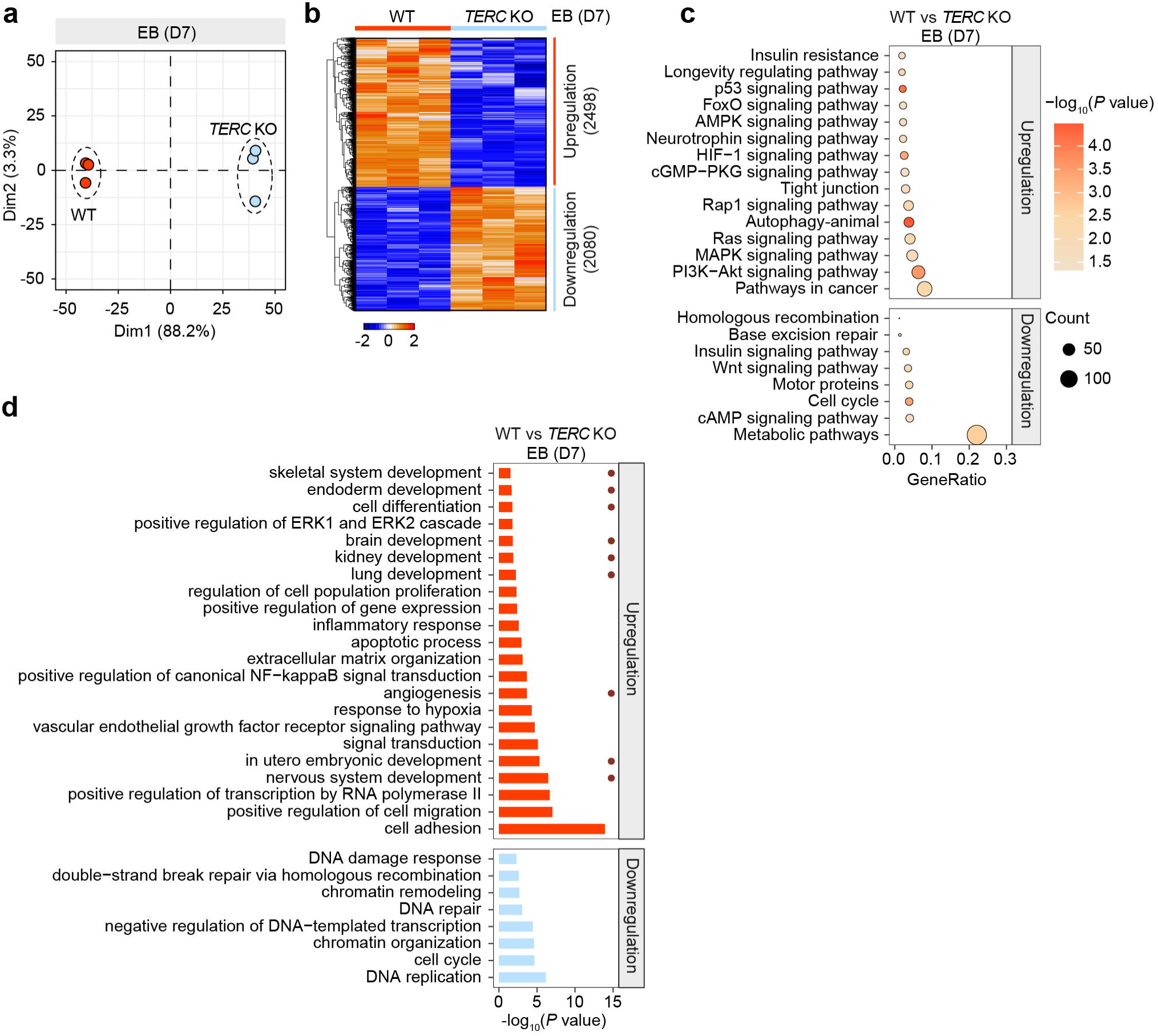
*TERC* deficiency disrupts developmental gene expression programs during differentiation. **a,** Principal component analysis of the transcriptomes of WT and *TERC* KO EBs on day 7 of *in vitro* differentiation. **b,** Heatmap of differentially expressed genes in WT vs. *TERC* KO EBs. The color key from blue to red indicates the relative gene expression level from low to high, respectively. **c,** KEGG pathway enrichment of differentially expressed genes in WT vs. *TERC* KO EBs. **d,** Significant functional enrichment of differentially expressed genes in WT vs. *TERC* KO EBs according to GO analysis of the RNA-seq data.

**Extended Data Fig. 4.**
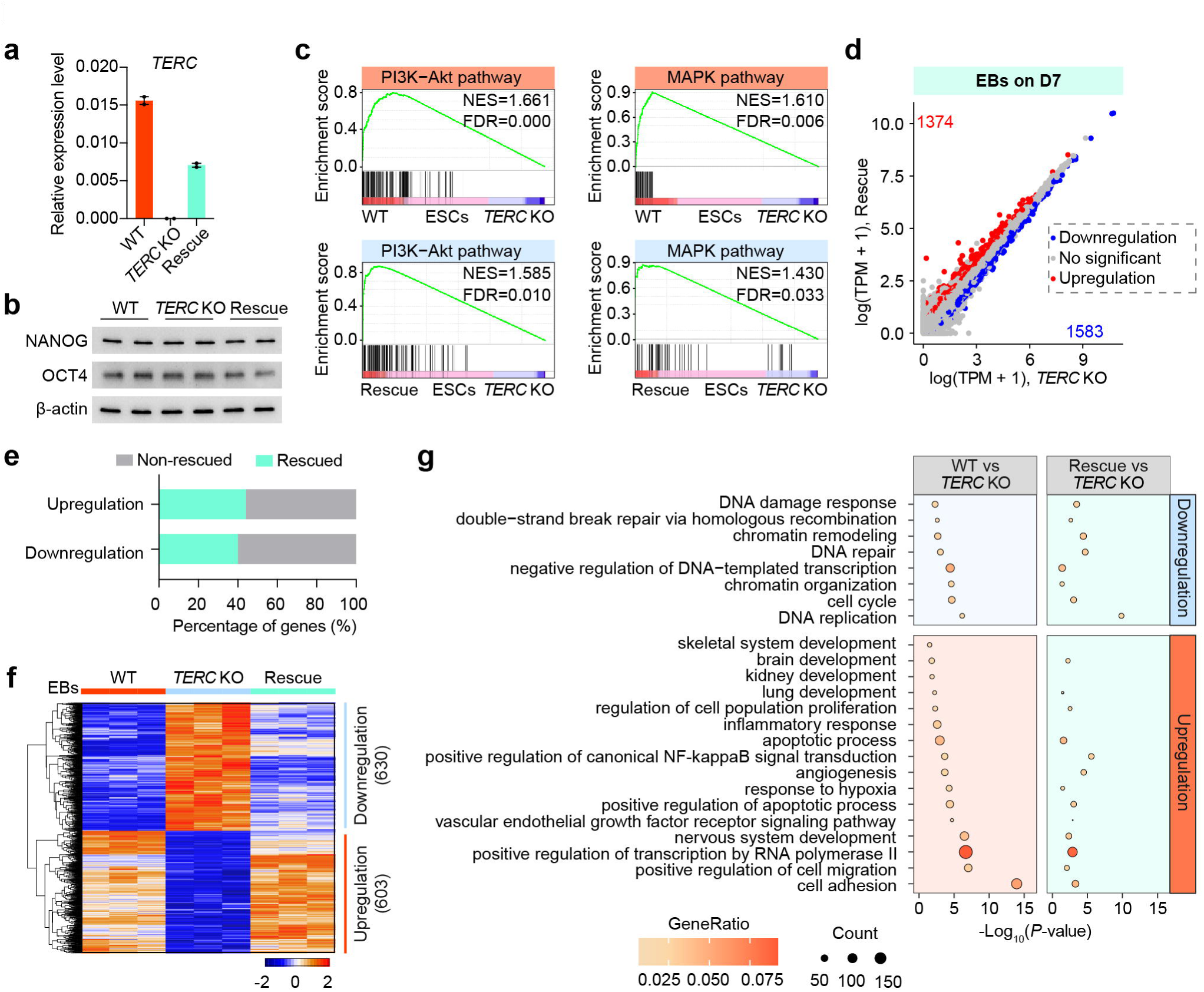
*TERC* re-expression rescues the differentiation of TERC KO hESCs *in vivo* and *in vitro*. **a,** qRT‒PCR analysis of the expression levels of *TERC* RNA in WT, *TERC* KO and TERC-rescue hESCs. The data are presented as the mean ± SEM; n=2 biological replicates. **b,** Western blot analysis of OCT4 and NANOG protein levels in WT, *TERC* KO and Rescue hESCs. β-actin was used as a loading control. **c,** GSEA showing enrichment of PI3K-Akt and MAPK pathway related genes in WT and *TERC* Rescue hESCs. **d,** Scatterplot showing the gene expression levels of *TERC* KO and *TERC* rescue EBs on day 7. The red and blue dots represent genes whose expression was upregulated and downregulated, respectively, in the *TERC* Rescue EBs compared with the *TERC* KO EBs. The gray dots represent genes whose expression did not significantly differ (no significant). **e,** Proportion of differentially expressed genes restored upon *TERC* re-expression. **f,** Heatmap depicting the expression profiles of rescued genes in EBs derived from WT, *TERC* KO and Rescue hESCs. **g,** GO analysis revealed significant functional enrichment of genes differentially expressed between WT/ *TERC* Rescue and *TERC* KO EBs.

**Extended Data Fig. 5.**
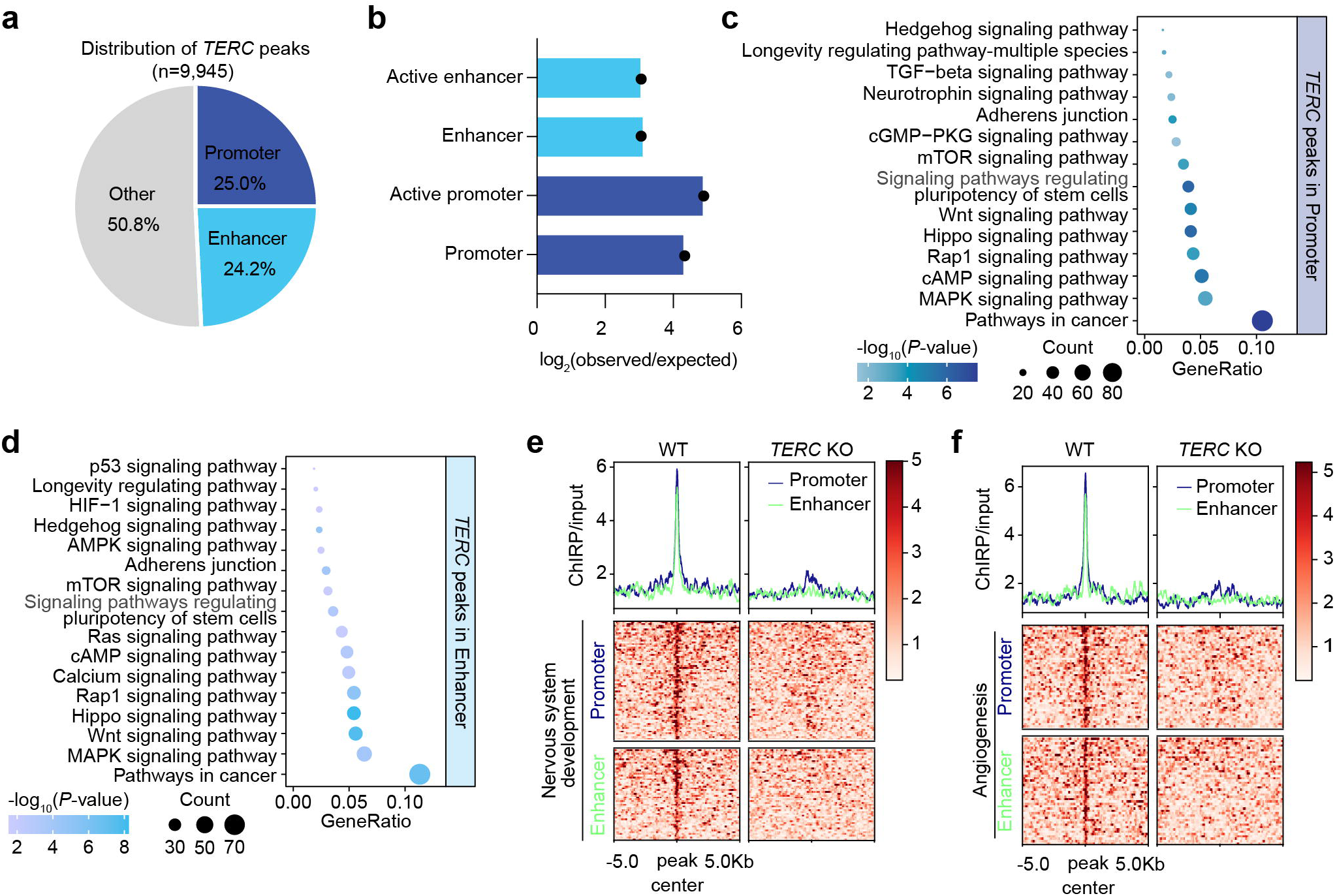
*TERC* binds to promoter and enhancer regions of differentiation and development genes. **a,** Pie chart showing regulatory element annotation of *TERC*-binding peaks. **b,** Fold enrichment of *TERC* peaks at each regulatory element by Genomic Association Test (GAT) analysis, and the log2(observed/expected) represents log2-transformed fold enrichment, given by the ratio of the observed count to the expected count. **c, d,** KEGG pathway enrichment for genes associated with *TERC*-bound promoters (**c**) and enhancers (**d**). The horizontal axis represents the gene ratio, the size of the circles represents the gene count, and the color depth represents the −log10-transformed *P* value. **e, f,** Heatmap showing enrichment of *TERC* at promoters and enhancers of genes associated with nervous system development and angiogenesis.

**Extended Data Fig. 6.**
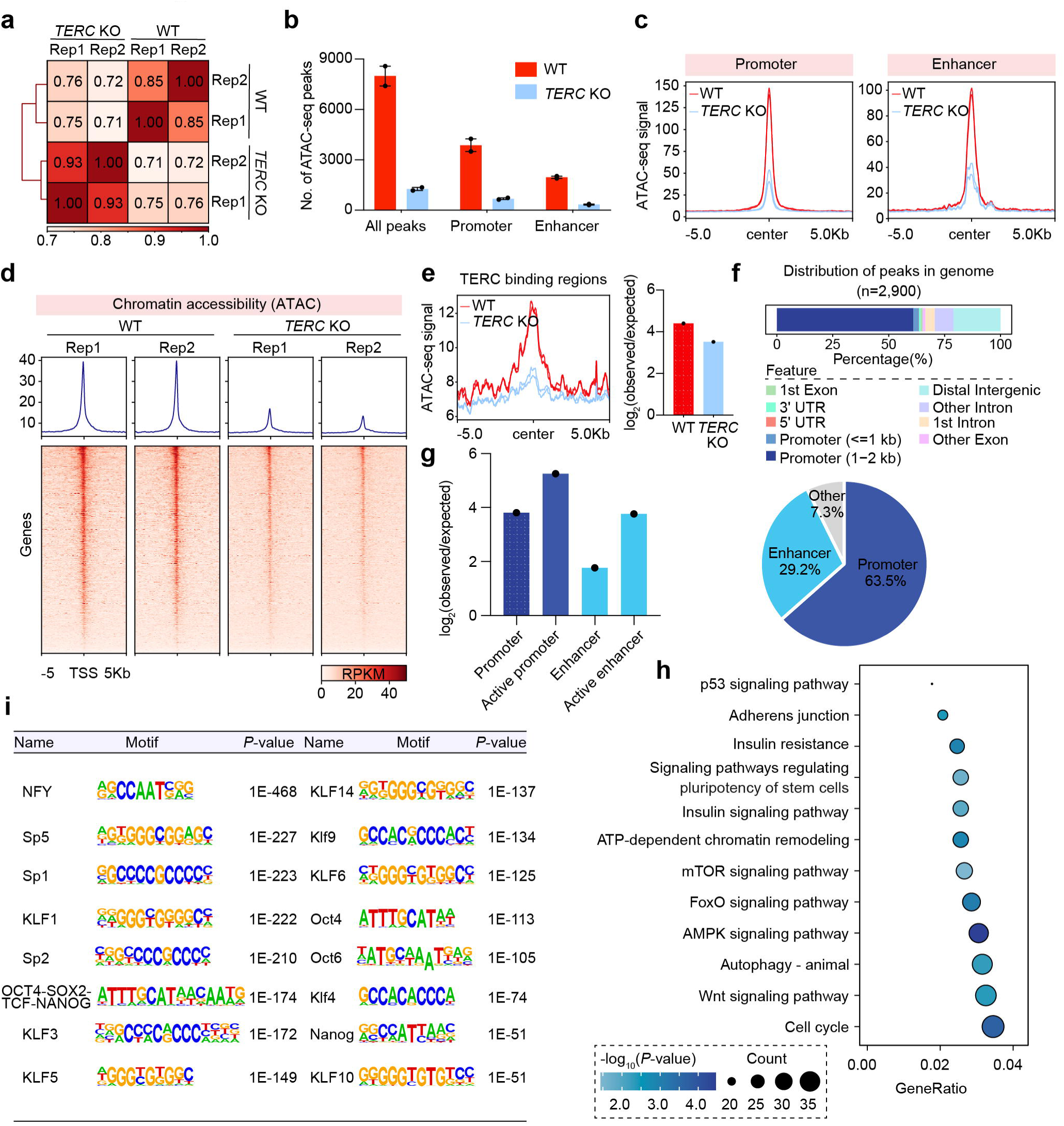
*TERC* promotes chromatin accessibility at differentiation and development genes. **a,** Scatterplots showing the genome-wide correlation of chromatin accessibility in WT and *TERC* KO hESCs. **b,** Bar plot showing the number of ATAC-seq peaks with significant enrichment (q value < 0.01) called by MACS2 in WT and *TERC* KO hESCs. Peaks annotated at promoters and enhancers are highlighted. **c,** Average profile of chromatin accessibility at promoters and enhancers in WT and *TERC* KO hESCs. **d,** Heatmap showing chromatin accessibility at ± 5 kb around the TSSs of RefSeq genes in WT and *TERC* KO hESCs. **e,** Changes in the chromatin accessibility of *TERC*-specific binding regions upon *TERC* knockout. Left: Accessibility profiles at *TERC* binding sites (n = 9,945). Right: GAT analysis showing the enrichment of ATAC-seq peaks at *TERC*-bound regions. **f,** Genomic distribution of downregulated ATAC-seq peaks in *TERC* KO hESCs compared with WT hESCs. **g,** Regulatory element annotation of downregulated ATAC-seq peaks in *TERC* KO hESCs compared with WT hESCs. The vertical axis represents the log2-transformed fold enrichment of each feature. **h,** Pathways enriched with genes marked by ATAC-seq peaks distributed in promoters and enhancers are significantly enriched, and these peaks are significantly less accessible in *TERC* KO hESCs than in WT hESCs. **i,** Key motifs identified from ATAC-seq peaks (n=2,900) that were significantly less accessible in *TERC* KO hESCs than in WT hESCs.

**Extended Data Fig. 7.**
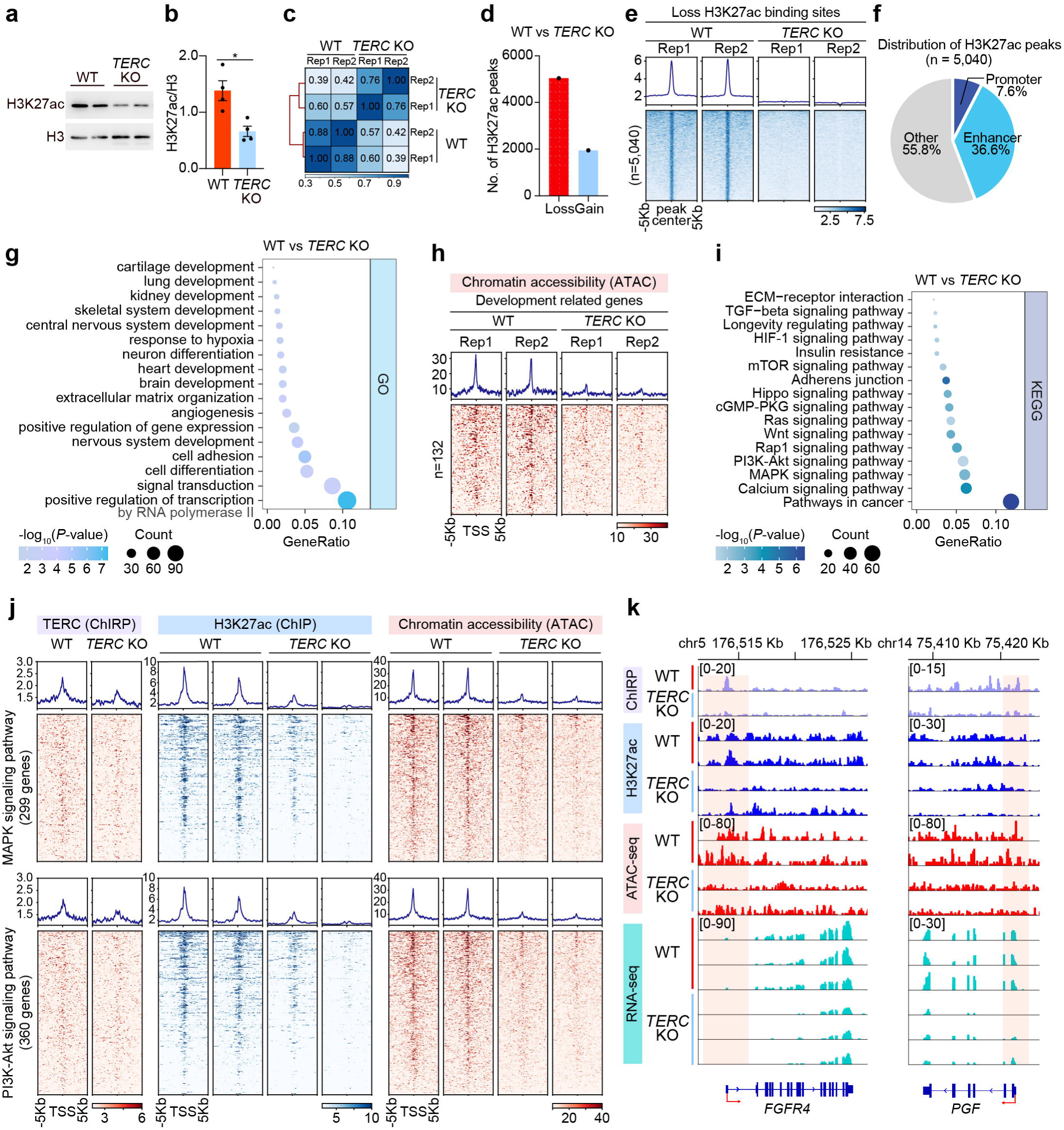
*TERC* depletion decreases H3K27ac occupancy at regulatory regions of developmental genes. **a, b,** Western blot analysis of H3K27ac protein levels in WT and *TERC* KO hESCs. H3 was used as a loading control, and *P* values were calculated via two-tailed unpaired t tests. **c,** Heatmap showing the correlation of H3K27ac occupancy between WT and *TERC* KO hESCs, with Pearson correlation coefficients indicated. Two biological replicates were performed for each genotype. **d,** Bar plot showing the number of H3K27ac peaks significantly lost or gained in hESCs upon *TERC* knockout. **e,** Heatmap showing H3K27ac occupancy at significantly lost H3K27ac peaks in *TERC* KO hESCs (n=5,040) compared with WT hESCs. **f,** Distribution of significantly lost H3K27ac peaks. **g,** Functional enrichment analyses for genes marked by H3K27ac peaks are shown in (**e**) and are distributed in promoter and enhancer regions (n=2,227). **h,** Heatmap showing ATAC-seq signals at the TSSs of development-related genes in WT and *TERC* KO hESCs. Development-related genes are derived from development-related enrichments, as shown in Figure (**g**). **i,** Significant pathways enriched with genes marked by H3K27ac peaks, as shown in (**e**), distributed in promoter and enhancer regions (n=2,227). **j,** Heatmap showing *TERC*, H3K27ac occupancy and ATAC-seq signals at the TSSs of genes associated with the MAPK and PI3K-Akt signalling pathways in WT and *TERC* KO hESCs. **k,** Genome browser representations of *TERC* ChIRP-seq, H3K27ac ChIP-seq, ATAC-seq and RNA-seq data at the *FGFR4* and *PGF* (MAPK or PI3K-AKT pathway) loci upon *TERC* knockout. The red arrow indicates the direction of transcription.

**Extended Data Fig. 8.**
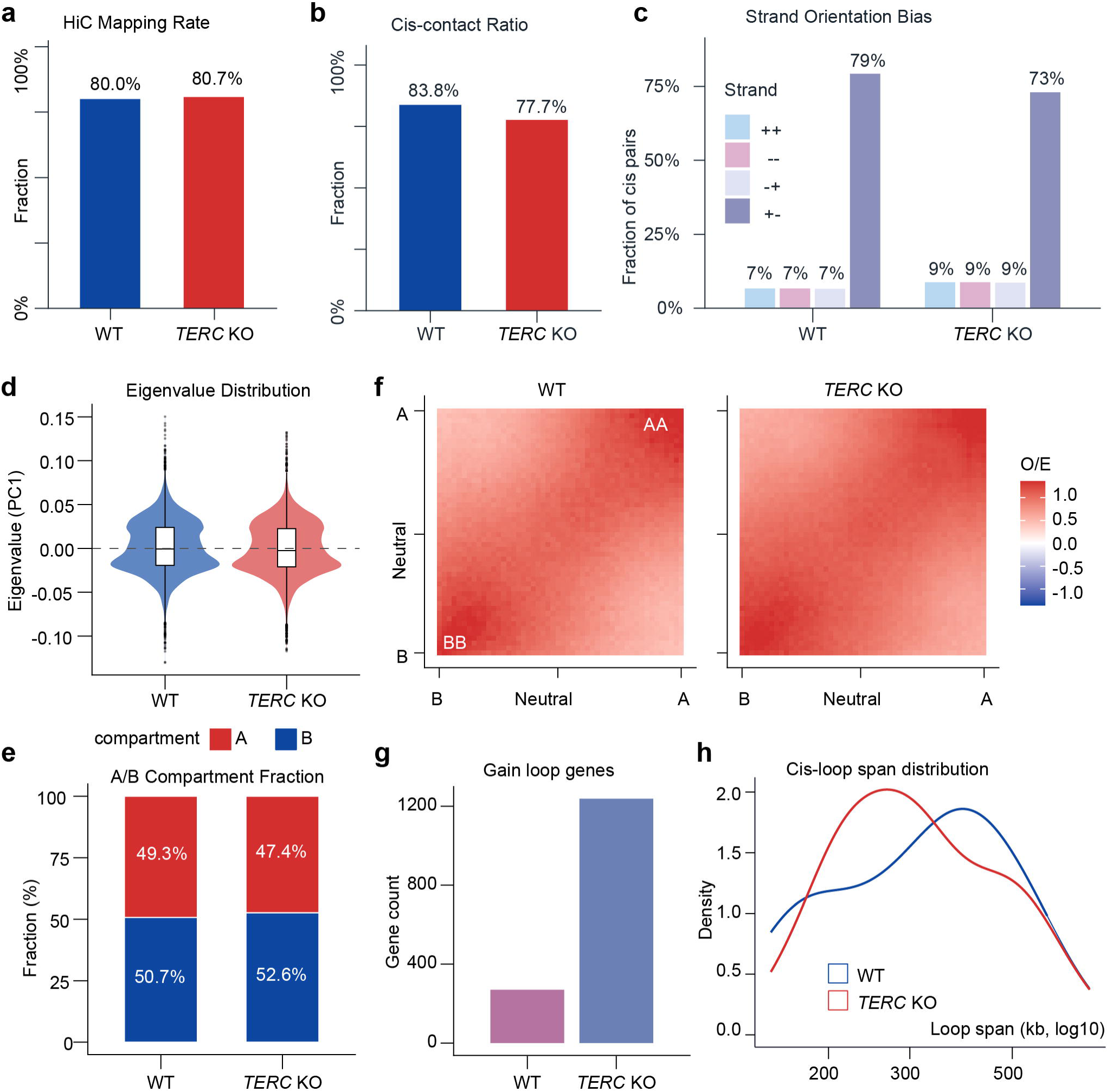
Hi-C library quality metrics, compartment distributions, saddle plots, and chromatin loop characteristics. **a,** Hi-C mapping rates for *TERC* KO (80.7%) and WT (80.0%) libraries, defined as the fraction of read pairs aligned to GRCh38/hg38 by BWA-MEM (v0.7.19). **b,** Cis-contact ratios for *TERC* KO (77.7%) and WT (83.8%), representing the fraction of valid (non-duplicate) pairs mapping to the same chromosome. **c,** Strand orientation bias of cis contact pairs for *TERC* KO and WT, shown as grouped bars for the four strand combinations (++, −−, −+, +−). The predominance of the +− orientation (*TERC* KO, 73%; WT, 79%) is characteristic of valid Hi-C proximity ligation products. **d,** Violin plots of the PC1 eigenvalue distribution from A/B compartment analysis at 100 kb resolution for WT (blue) and *TERC* KO (red) cells, with embedded box plots indicating the median and interquartile range. The bimodal distribution reflects the separation of A (positive) and B (negative) compartments. **e,** Stacked bar charts showing the genome-wide A/B compartment fractions in WT (A: 49.3%; B: 50.7%) and *TERC* KO (A: 47.4%; B: 52.6%) cells. *TERC* knockout results in a modest increase in B compartment occupancy (+1.9%). **f,** Saddle plots of observed-over-expected (O/E) contact enrichment (log₂ scale) as a function of the eigenvector quantile for WT (left) and *TERC* KO (right) cells. Bins are ranked across 50 quantiles from B (bottom-left) to A (top-right). The AA (top-right) and BB (bottom-left) corners represent homotypic compartment self-interactions. O/E values were computed using cooltools. **g,** Bar chart comparing the number of genes associated with gained chromatin loops in WT versus *TERC* KO. *TERC* KO hESCs exhibit a substantially greater number of genes with gained loop anchors, indicating widespread de novo chromatin loop formation. **h,** Kernel density estimates of the cis-loop genomic span (log₁₀ scale, kb) for WT (blue) and *TERC* KO (red). Loops were called at 15 kb resolution using Mustache (v1.3.3; *P* < 0.05; maximum span: 2 Mb). The leftward shift of the *TERC* KO distribution indicates a predominance of shorter-range interactions in *TERC* KO relative to the WT cells.

**Extended Data Fig. 9.**
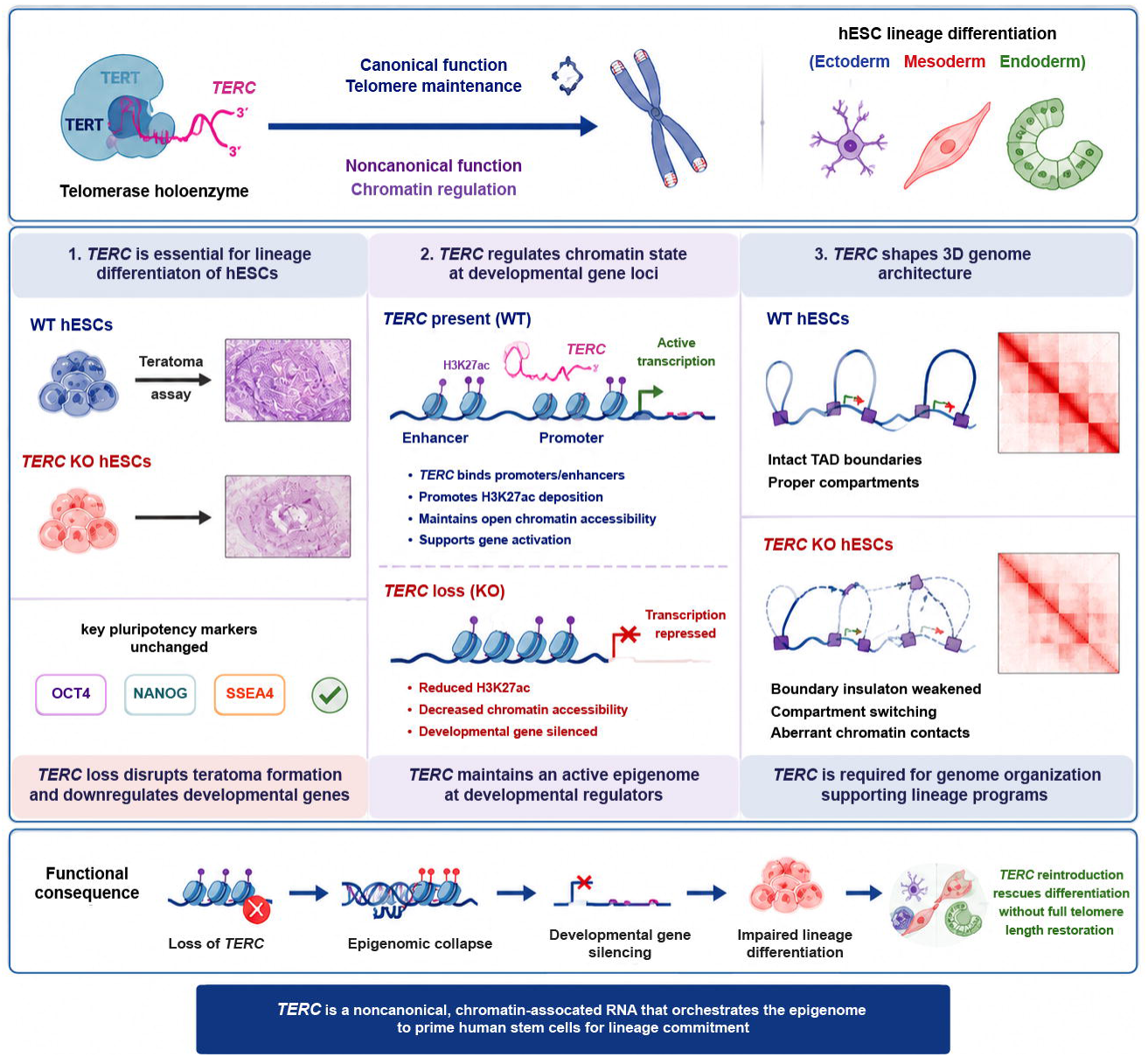
A schematic diagram illustrating the role of telomerase RNA in mediating chromatin accessibility for lineage commitment.

**Supplementary Table 1.**
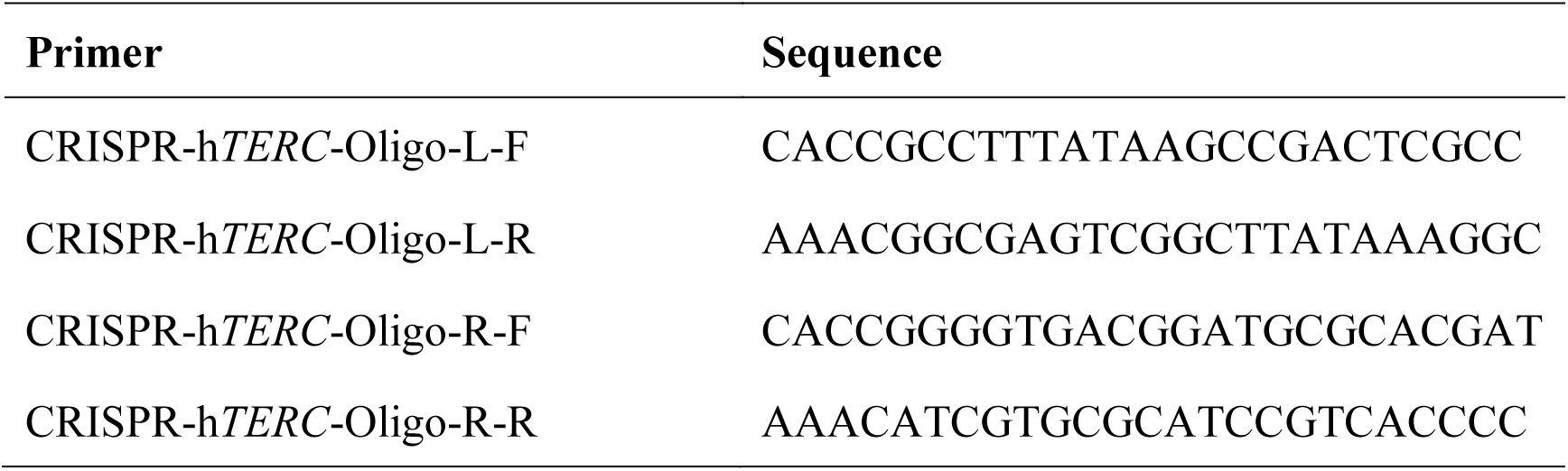
Primers for CRISPR/Cas9 experiments.

**Supplementary Table 2.**
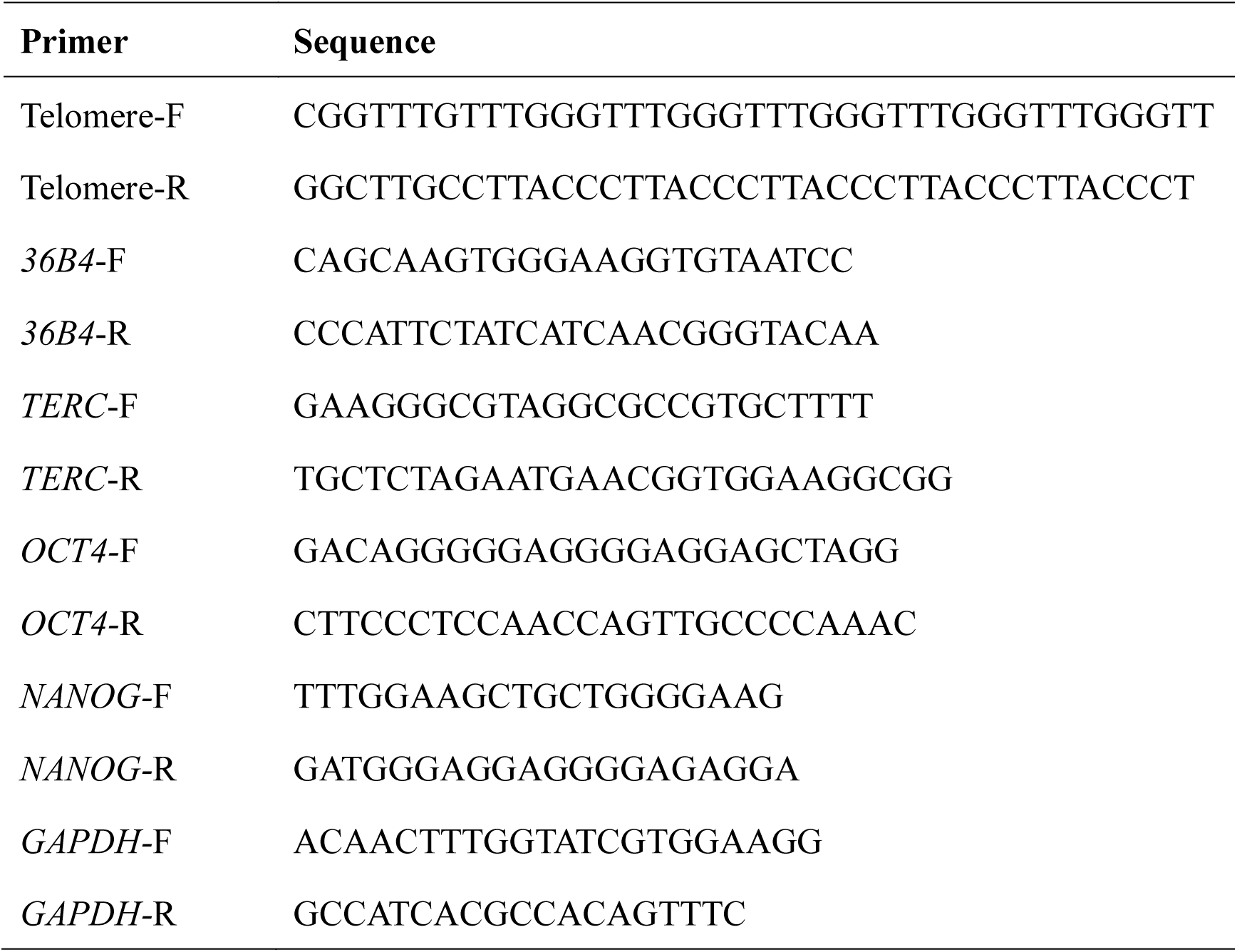
Primers for the T/S ratio and RT‒qPCR.

